# Deciphering the role of histone modifications in memory and exhausted CD8 T cells

**DOI:** 10.1101/2025.04.16.649198

**Authors:** Hua Huang, Amy E. Baxter, Zhen Zhang, Charly R. Good, Katherine A. Alexander, Zeyu Chen, Paula A. Agudelo Garcia, Parisa Samareh, Sierra M. Collins, Karl M. Glastad, Lu Wang, Gregory Donahue, Sasikanth Manne, Josephine R. Giles, Junwei Shi, Shelley L. Berger, E. John Wherry

**Author notes:** Address correspondence to: S.L.B. and E.J.W. These authors contributed equally (Hua Huang, Amy E. Baxter, Zhen Zhang, Charly R. Good, Katherine A. Alexander).

## Abstract

Exhausted CD8 T cells (T_EX_) arising during chronic infections and cancer have reduced functional capacity and limited fate flexibility that prevents optimal disease control and response to immunotherapies. Compared to memory (T_MEM_) cells, T_EX_ have a unique open chromatin landscape underlying a distinct gene expression program. How T_EX_ transcriptional and epigenetic landscapes are regulated through histone post-translational modifications (hPTMs) remains unclear. Here, we profiled key activating (H3K27ac and H3K4me3) and repressive (H3K27me3 and H3K9me3) histone modifications in naive CD8 T cells (T_N_), T_MEM_ and T_EX_. We identified H3K27ac-associated super-enhancers that distinguish T_N_, T_MEM_ and T_EX_, along with key transcription factor networks predicted to regulate these different transcriptional landscapes. Promoters of some key genes were poised in T_N_, but activated in T_MEM_ or T_EX_ whereas other genes poised in T_N_ were repressed in T_MEM_ or T_EX_, indicating that both repression and activation of poised genes may enforce these distinct cell states. Moreover, narrow peaks of repressive H3K9me3 were associated with increased gene expression in T_EX_, suggesting an atypical role for this modification. These data indicate that beyond chromatin accessibility, hPTMs differentially regulate specific gene expression programs of T_EX_ compared to T_MEM_ through both activating and repressive pathways.

## INTRODUCTION

CD8 T cells are a proliferative and functional differentiation hierarchy. Following activation, quiescent naive CD8 T cells (T_N_) undergo a complex and extensive rewiring of epigenetic, transcriptional regulation and gene expression programs. In the days after activation, T_N_ differentiate into two major divergent populations: short-lived cytotoxic effector CD8 T cells (T_EFF_) and the precursors for long-lived, quiescent memory CD8 T cells (T_MEM_)^1,2^. Whereas T_EFF_ mediate antiviral and anticancer effector functions, migrate throughout the body and help control initial disease, T_MEM_ precursors are more restrained. Following antigen clearance, most T_EFF_ die, but T_MEM_ precursors survive and differentiate into mature T_MEM_ that are quiescent and slowly self-renew. Furthermore, T_MEM_ can rapidly reactivate effector functions and proliferate following encounter with the same antigen, mounting robust recall responses. However, if antigen persists, such as during chronic viral infections and cancer, these early precursors instead differentiate into exhausted CD8 T cells (T_EX_). In contrast to T_MEM_, T_EX_ are maintained by persistent stimulation, resulting in chronic activation and altered effector capacity. Thus, T_EX_ mount weak recall responses to antigen restimulation and are associated with poor disease control compared to T_MEM_^3^. Despite both cell populations originating from a common precursor cell (T_N_), T_MEM_ and T_EX_ represent divergent differentiation paths, resulting in highly distinct cell types. This ability of a common progenitor population to give rise to differentiation hierarchies consisting of diverse cell types with distinct functions has parallels throughout developmental biology, for example in the gastrointestinal tract where an intestinal stem cell niche gives rise to all mucosal cell types^4^. Although much work has focused on the functional differences between T_MEM_ and T_EX_, the precise epigenetic and transcriptional mechanisms associated with differentiation of these divergent cell states remains to be fully defined.

Epigenetic profiling of chromatin accessibility by Assay for Transposase-Accessible Chromatin with sequencing (ATAC-seq) has revealed that T_MEM_ and T_EX_ are as epigenetically distinct from each other as each cell type is from T_N_^5–10^. Furthermore, T_EX_ have a unique chromatin accessibility profile with distinct open chromatin sites compared to other CD8 T cell subsets^5,6^. These observations support the idea that T_MEM_ and T_EX_ are separate lineages of mature CD8 T cells. The unique chromatin accessibility landscape of T_EX_ is established in part by the thymocyte selection associated high mobility group transcription factor (TF) TOX^11–16^. TOX is essential for the formation of T_EX_ but is dispensable for the generation of T_MEM_ despite transient expression of TOX following acute stimulation^13,17^. In addition, TOX may regulate further differentiation once the T_EX_ population is established^18^, coordinating transitions between progenitor, intermediate and terminal T_EX_ subsets^19–25,18^. Thus, TOX plays a key role in establishing the unique T_EX_ open chromatin landscape profiled by ATAC-seq. However, the associations between this T_EX_ open chromatin landscape and regulation of gene expression through histone post-translational modifications (hPTMs), including how these hPTMs change as T_N_ differentiate into T_MEM_ or T_EX_, remains incompletely understood.

Indeed, CD8 T cell differentiation is regulated by the addition or removal of hPTMs by a variety of epigenetic enzymes. In all multicellular organisms, histone 3 lysine 27 acetylation (H3K27ac) is associated with active enhancers, whereas methylation at the same site (H3K27me3) is associated with decreased gene expression and the formation of facultative heterochromatin. In CD8 T cells, EZH2, a histone methyltransferase that establishes H3K27me3, and KDM6B, a lysine demethylase that removes methyl groups from H3K27, have both been reported to regulate cell fate specification between T_EFF_ and T_MEM_ populations, as well as formation of T_MEM_ capable of mounting robust recall responses^26–28^. Furthermore, the histone deacetylase HDAC3, which leads to chromatin compaction in part by removing acetyl groups from H3K27ac, restrains T_EFF_ development and dampens effector function^29^. In addition to H3K27me3, H3K9me3 is also associated with heterochromatin and regulates constitutive repression of repeated DNA elements and long-term repression of inactive regions. The methyltransferase Suv39h1 that establishes H3K9 tri-methylation silences T_MEM_-associated genes to enable T_EFF_ differentiation, and may regulate T_EX_ effector function^30,31^. Furthermore, epigenetic enzymes such as protein arginine methyltransferase PRMT4 (CARM1)^32^, chromatin remodelers SWI/SNF family members BAF and PBAF^33–37^, and ASXL1^38^, as well as enzymes catalyzing DNA methylation and demethylation (TET2^39,40^ and DNMT3A^41–43^) have all been implicated in regulating fate transitions between and/or within CD8 T cell subsets. The diverse functions of the epigenetic enzymes that potentially regulate CD8 T cell differentiation suggests that complex patterns of hPTMs may be a feature of the distinct T_N_, T_MEM_ and T_EX_ epigenetic landscapes. Interrogating the differences in these hPTM patterns may provide insights into the diverse transcriptional regulation and gene expression programs of these CD8 T cell states.

Once established, T_EX_ are fate inflexible and do not differentiate into T_EFF_ or T_MEM_ cells. Furthermore, T_EX_ retain the epigenetic “scars” of exhaustion even after removal of antigen and “cure” of chronic infection^44^. In contrast, T_MEM_ are fate-flexible and are poised to rapidly respond when re-encountering antigen by differentiating into highly functional T_EFF_. This T_EX_ fate inflexibility limits disease control. Targeted immunotherapies such as PD-1 pathway blockade “reinvigorate” T_EX_ and have revolutionized cancer therapy^45–47^. However, not all patients experience clinical benefit, in part because the burst of effector activity in reinvigorated T_EX_ is transient. This inability to provoke durable changes in T_EX_ function and differentiation is due, at least in part, to the failure of T_EX_-targeted immunotherapies to rewire the T_EX_ chromatin landscape and epigenetically reprogram these cells into T_EFF_ or T_MEM_ cells^5,6,18^. Indeed, because PD-1 pathway blockade does not change the T_EX_ open chromatin landscape^5,10^, reinvigorated T_EX_ revert to their original exhausted state over time^5^. Thus, although the open chromatin landscape of T_EX_ has been defined by ATAC-seq, a more comprehensive understanding of how changes in chromatin accessibility and hPTMs are associated with gene expression is required to develop immunotherapy strategies that provoke durable responses in T_EX_. Furthermore, how epigenetic regulation through combinations of hPTMs might regulate the maintenance of T_EX_ epigenetic scars, but enable fate flexibility in T_MEM_, remains unclear.

To address these questions, we interrogated hPTMs patterns in T_N_, T_MEM_ and T_EX_ to investigate how active and repressive hPTMs were associated with the diverse gene expression and chromatin accessibility landscapes of these distinct CD8 T cell differentiation states. The epigenetic transition from a quiescent T_N_ state into antigen-experienced populations (i.e. from T_N_ to T_MEM_ or T_EX_) was associated with broad genome-wide alterations, highlighting the extensive epigenetic remodeling associated with CD8 T cell activation. Moreover, these genome-wide changes were associated with distinct chromatin features for T_MEM_ and T_EX_ cells, highlighting a key role for hPTM patterns in the development of fate-flexible T_MEM_ versus fate-inflexible T_EX_. Defining these hPTM patterns may provide insight into the control of gene expression in T cell populations with diverse functions. These data provide a foundation for future epigenetic-based therapeutic approaches.

## RESULTS

### Histone modifications are associated with distinct gene expression landscapes of T_MEM_ and T_EX_

To study how hPTMs could regulate T_EX_ development, we profiled activating and repressive histone modifications in T_EX_ compared to T_MEM_ and T_N_. We used different strains of lymphocytic choriomeningitis virus (LCMV) to induce either an acutely resolving infection [Armstrong (Arm)] with development of T_EFF_ followed by T_MEM_, or to establish a chronic infection [clone13 (Cl13)] that results in T_EX_ formation^3^. We adoptively transferred a physiological number of naive T cell receptor (TCR) transgenic LCMV D^b^GP33-41-specific CD8 T cells (P14 cells) into congenically distinct recipient mice and infected these recipient mice with LCMV Arm or Cl13 (Fig. 1a; ^48,49^). At ∼day 30 post-infection, we isolated P14 cells from mice infected with either Arm (T_MEM_, Fig. S1a-S1c) or Cl13 (T_EX_, Fig. S1d-S1f). P14 cells from an uninfected mouse were isolated as naive controls (T_N_). We then performed Cleavage Under Targets and Release Using Nuclease (CUT&RUN) for histone modifications H3K27ac, H3K4me3, H3K27me3, and H3K9me3 on T_N_, T_MEM_, and T_EX_ cells, alongside RNA-sequencing (Fig. 1a).

**Figure 1.**
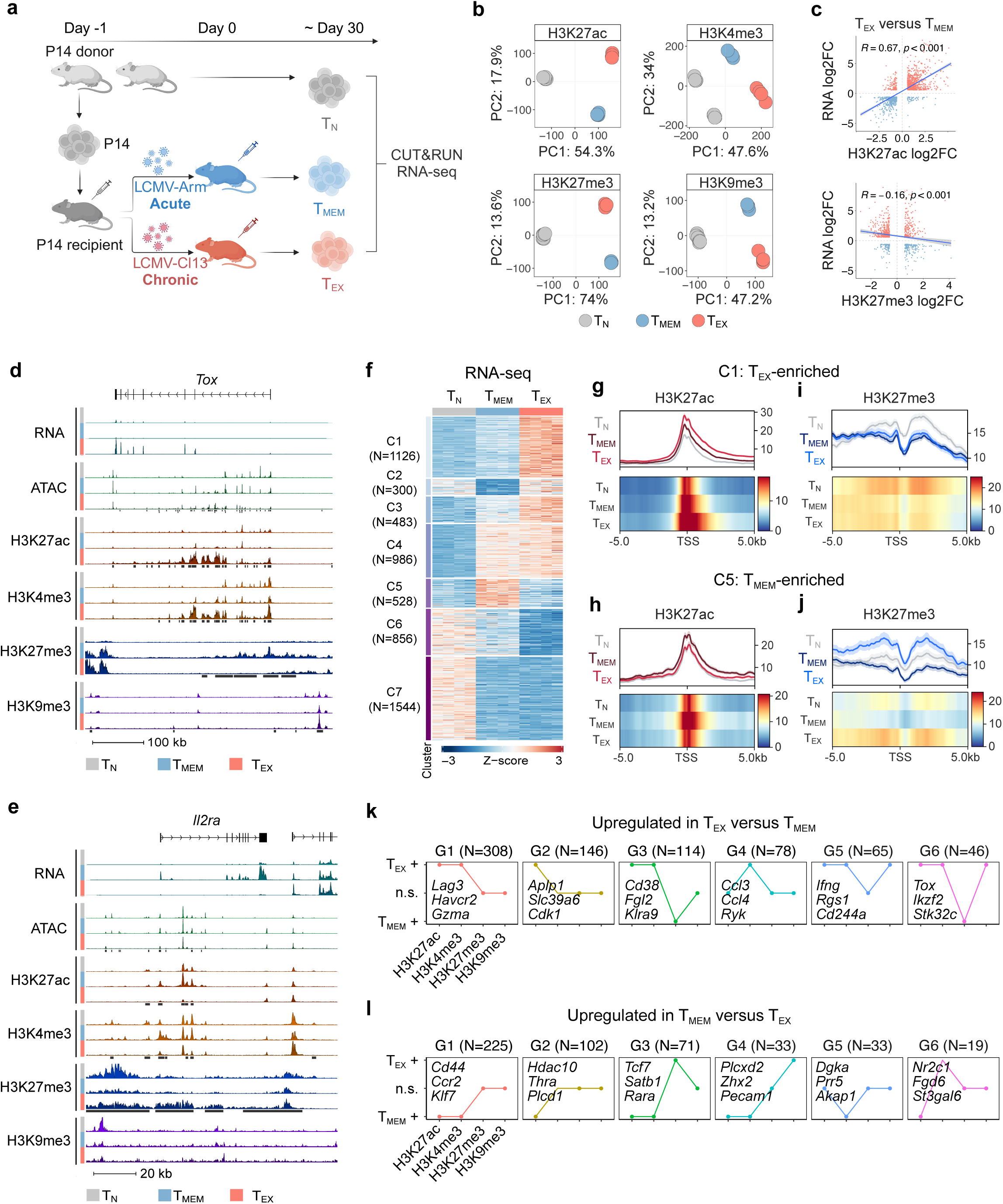
**Histone modifications act in concert to regulate gene expression in T_EX_ and T_MEM_** (**a**) Experimental design. (**b**) PCA of H3K27ac, H3K4me3, H3K27me3 and H3K9me3 data for T_N_, T_MEM_ and T_EX_. For T_MEM_ and T_EX_, n = 4 biological replicates; for T_N_, n = 8 biological replicates. (**c**) Correlation of change in RNA expression between T_MEM_ and T_EX_ and change in H3K27ac (top), or H3K27me3 (bottom). R and associated P-value represent Pearson correlation. Each dot represents one gene with one associated peak. (**d-e**) Genome tracks showing RNA-seq, ATAC-seq and hPTM data. Differentially modified regions between T_MEM_ and T_EX_ for each hPTM are highlighted in black boxes. (**f**) Heatmap of DEGs showing K-mean clusters for all pairwise comparisons between T_N_, T_MEM_, and T_EX_. (**g-h**) Meta plot (top) and heatmap plot (bottom) of H3K27ac at TSS for DEGs **(g)** cluster 1 (C1) and (**h**) cluster 5 (C5). (**i-j**) Meta plot (top) and heatmap plot (bottom) of H3K27me3 at TSS for DEGs **(i)** cluster 1 (C1) and **(j)** cluster 1 (C5). (**k-l**) Comparison of hPTM patterns between T_MEM_ and T_EX_ for **(k)** genes with increased expression in T_EX_ compared to T_MEM_ or **(l)** genes with increased expression in T_MEM_ compared to T_EX_. Top six most frequent groups for each set of genes plotted.

We first investigated whether hPTM patterns were distinct between CD8 T cells from acute and chronic infection compared to naive control cells. Principal component analysis (PCA) revealed that T_N_, T_MEM_ and T_EX_ occupied separate regions of PCA space for each hPTM studied (Fig. 1b). Furthermore, for all hPTMs, T_N_ clustered separately from T_EX_ and T_MEM_ in PC1 whereas T_MEM_ and T_EX_ separated in PC2 (Fig. 1b). Thus, the highest numbers of differential hPTMs were identified between T_N_ and either T_MEM_ or T_EX_ (Fig. S1g), suggesting that the greatest magnitude of hPTM changes occurred as T_N_ were activated and differentiated into T_EX_ or T_MEM_ cell fates.

To examine how hPTMs were associated with changes in gene expression between the three CD8 T cell subtypes, we first mapped each hPTM region to the nearest gene. We selected the peak for each gene that was the most variable across conditions, mapping one peak per gene. We then correlated the fold change in RNA expression of these genes to the fold change in hPTMs at the gene-associated peak. Genes with higher H3K27ac were associated with increased gene expression (R = 0.67, Fig. 1c, top and Fig. S1h-S1i). In contrast, higher H3K27me3 was only weakly correlated with lower gene expression (R = -0.16, Fig. 1c, bottom and Fig. S1h-S1i). However, key CD8 T cell genes had concurrent changes in H3K27ac and H3K27me3. For example, the TF *Tcf7* was highly expressed in T_N_, moderately expressed in T_MEM_ but lower in T_EX_ (Fig. S1j, upper tracks). These differences in RNA expression were associated with concordant changes in activating hPTMs: H3K27ac was highest in T_N_, moderate in T_MEM_ and minimal in T_EX_. In contrast, the repressive modification H3K27me3 was low in both T_N_ and T_MEM_, but higher in T_EX_ (Fig. S1j, lower tracks). Thus, reduced *Tcf7* expression in T_MEM_ compared to T_N_ was associated with a decrease in activating hPTMs, whereas in T_EX_ this gene had a combination of both lower H3K27ac and higher H3K27me3. This combination of changes was associated with the lowest *Tcf7* RNA expression between CD8 T cell subsets.

The exhaustion-associated TF *Tox* is highly expressed in T_EX_ compared to T_MEM_ and the *Tox* locus has extensive open intronic chromatin in T_EX_^13^ (Fig. 1d). This region of the *Tox* gene was extensively marked with both activating H3K27ac and H3K4me3 in T_EX_, concurrent with low H3K27me3 (Fig. 1d, Fig. S1k). In contrast, reduced expression of IL-2 receptor alpha (*Il2ra*) in T_EX_ compared to T_MEM_ was associated with higher H3K27me3 and lower H3K27ac and H3K4me3 (Fig. 1e, Fig. S1k). These analyses suggest that combinatorial changes of activating and repressive hPTMs accompany changes in gene expression between CD8 T cell fates.

To probe how the interplay between different hPTMs might regulate gene expression between CD8 T cell states, we next focused on hPTMs at promoters. H3K27ac at promoters is associated with active transcription, whereas H3K27me3 is present at inactive or poised promoters^50^. K-mean clustering identified 7 groups of genes differentially expressed between T_N_, T_EX_ and T_MEM_ (Fig. 1f). One set of differentially expressed genes (DEGs) was highly expressed in T_EX_ compared to both T_N_ and T_MEM_ (Fig. 1f: C1; 1126 genes), whereas a second set of DEGs was highly expressed in T_MEM_ compared to both T_N_ and T_EX_ (Fig. 1f: C5; 528 genes). For both clusters of DEGs, H3K27ac levels were highest and broadest around the transcription start site (TSS) in the corresponding CD8 T cell type (Fig. 1g-1h). For example, for genes upregulated in T_EX_ (C1), H3K27ac peaks at the TSS were highest and broadest in T_EX_ (Fig. 1g), whereas C5 DEGs (upregulated in T_MEM_) had higher H3K27ac in T_MEM_ than T_EX_ and T_N_ (Fig. 1h). In contrast, the association between H3K27me3 and gene expression varied between cell state-associated genes. H3K27me3 was lower at the TSS in both T_EX_ and T_MEM_ compared to T_N_ for C1 genes, despite elevated gene expression for this cluster only in T_EX_ (Fig. 1i). However, H3K27me3 was lowest at the TSS of C5 genes (upregulated in T_MEM_) in T_MEM_ compared to T_N_ and T_EX_ (Fig. 1j), suggesting that loss of this repressive mark may have a distinct role in enforcing gene expression in T_MEM_. Furthermore, H3K27me3 was higher at the TSS of C5 genes in T_EX_ compared to both T_MEM_ and T_N_ (Fig. 1j) provoking the hypothesis that a subset of genes expressed in T_MEM_ are actively repressed in T_EX_. Finally, consistent with the role of silencing hPTMs in gene repression, H3K27me3 deposition was distributed over ∼10 kb around the TSS (Fig. 1j). These findings further support a role for changes in both activating and repressive hPTMs in regulating gene expression in T_MEM_ and T_EX_, including at and outside the promoter, and also indicate that gain of activation marks, rather than loss of repressive modifications, may be a more common feature associated with increased gene expression.

We next investigated the genome-wide association of combinatorial changes in hPTMs with gene expression. DEGs between T_EX_ and T_MEM_ were binned into patterns based on higher or lower abundance of the hPTMs analyzed. For genes upregulated in T_EX_ compared to T_MEM_, 5 of the 6 most frequent patterns were characterized by increased H3K27ac deposition in T_EX_ (Fig. 1k; G1, G2, G3, G5, and G6), which commonly co-occurred with higher H3K4me3 in T_EX_ (Fig. 1k; G1, G3, G5, G6). In contrast, associations between T_EX_ gene expression and repressive marks were less consistent. H3K27me3 and H3K9me3 levels were unchanged in T_EX_ versus T_MEM_ in 3 of the 6 patterns (G1, G2 and G4), with a shift to lower H3K27me3 in T_EX_ only in patterns G3 and G6. Instead, H3K9me3 levels increased concurrently with H3K27ac and H3K4me3 in 2 patterns (G5 and G6), including for *Tox* (G6). Similar features were identified for genes upregulated in T_MEM_ (Fig. 1l). Indeed, the top 3 most frequent hPTM patterns were similar between genes upregulated in T_MEM_ or T_EX_ (Fig. 1k-1l; G1, G2, G3). Together, these analyses suggest that gain of H3K27ac is the most prevalent and homogenous feature of differential gene expression between T_EX_ and T_MEM_. In contrast, loss of the H3K27me3 is linked to increased expression of only a subset of genes whereas H3K9me3 may be associated with increased expression of some genes in T_EX_.

### H3K27ac identifies putative enhancers and predicts families of transcription factors acting at these enhancers in T_EX_

Analysis of TF binding sites in differentially accessible chromatin regions has identified TFs driving distinct T_MEM_ and T_EX_ cell fates^10,51^. However, chromatin accessibility defined by ATAC-seq alone may not be sufficient to identify functional enhancers. Therefore, analysis of TF activity at total accessible chromatin sites may not reflect the TF regulatory networks active at enhancers controlling gene regulation. As H3K27ac amplitude is strongly correlated with enhancer activity^52,53^, we hypothesized that identifying TF activity at H3K27ac sites could predict TFs active at putative enhancers. Thus, we performed Taiji PageRank analysis^54^ on H3K27ac data to identify TF networks at putative enhancers that might drive T_EX_ differentiation. Taiji PageRank analysis combines peak intensity, TF motif binding site accessibility, and TF expression to predict TF activity. TFs associated with quiescence such as LEF1 and TCF7 ranked highly in T_N_ and higher in T_MEM_ than in T_EX_ (Fig. 2a)^55,56^, whereas TFs reported to coordinate memory versus effector responses, including RUNX1 and RUNX2, ranked highest in T_MEM_^57^ (Fig. 2a). BATF, NFAT and IRF family members (IRF1, IRF3, IRF4 and IRF8) ranked highly in T_EX_ (Fig. 2a), supporting identified roles for these TFs in exhaustion^58–62^. To identify TFs with predominant roles at putative enhancers (i.e. H3K27ac sites) rather than all open chromatin regions, we compared Taiji PageRank scores generated using H3K27ac (Fig. 2a) to published ATAC-seq data (Fig. S2a)^5^. The majority of TFs that ranked highly in T_EX_ compared to T_MEM_ using H3K27ac also ranked highly in ATAC-seq-based analysis, including NFAT, AP-1 and BATF (Fig. 2b). Correspondingly, TFs that ranked more highly in T_MEM_ compared to T_EX_ in the H3K27ac analysis also ranked highly using ATAC-seq (e.g. TCF7) (Fig. 2b). However, some TFs were differentially ranked between the two analyses. Whereas NUR77 (NR4A1) was highly ranked in T_EX_ compared to T_MEM_ based on chromatin accessibility^63,64^, this TF had comparable rankings in T_EX_ and T_MEM_ when assessed using enhancer-biased H3K27ac sites (Fig. 2b). In contrast, ZEB1 scored highly using H3K27ac in T_EX_ compared to T_MEM_, but had similar PageRank scores in T_MEM_ and T_EX_ by ATAC-seq (Fig. 2b). The ZEB family of TFs, ZEB1 and ZEB2, may have reciprocal and potentially opposing roles in mature CD8 T cell differentiation. Of note, the ZEB2 binding motif remains undefined, preventing Taiji PageRank analysis. Whereas ZEB1 is required for T_MEM_ survival and function^10,65^, ZEB2 promotes terminal differentiation and effector function^66,67^. In T_EX_, ZEB1 regulates T_EX_ survival and persistence, whereas ZEB2 mediates T_EX_ cytotoxic function, suggesting these TFs modulate distinct, potentially opposing, gene regulatory networks^10,68^. A deeper understanding of TFs acting at putative enhancers may uncover relevant T_EX_ transcriptional networks, including transcriptional pathways regulated by the TF pair ZEB1/ZEB2.

**Figure 2.**
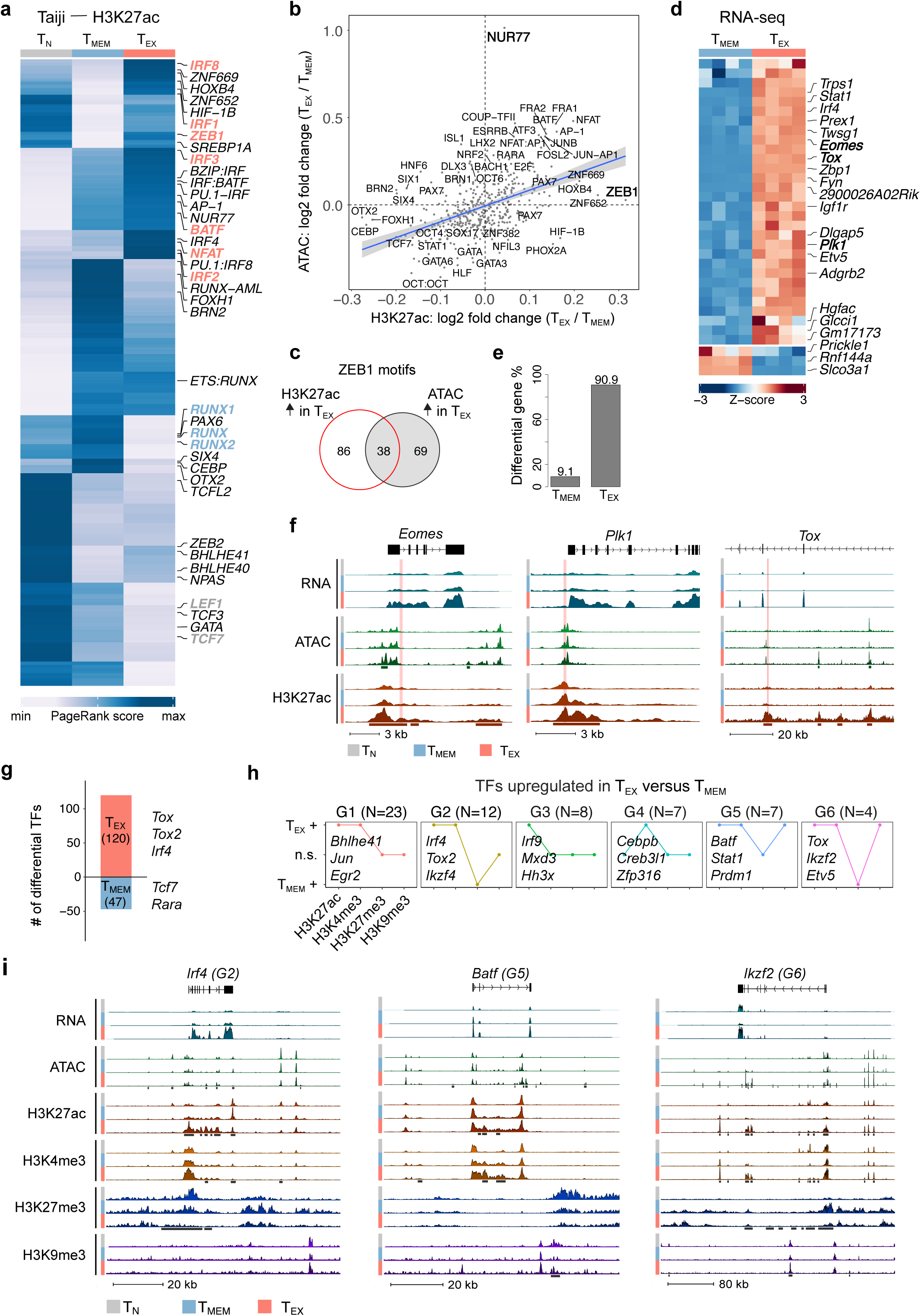
**Identification of predicted TF binding motifs under H3K27ac identities role for ZEB1 in T_EX_** (**a**) Heatmap of normalized Taiji PageRank scores determined using RNA-seq and H3K27ac data. (**b**) Correlation plot comparing Taiji PageRank scores from H3K27ac to ATAC-seq data. Axes represent log2 fold change (log2FC) in Taiji PageRank scores between T_EX_ and T_MEM_ (**c**) Venn diagram comparing number of ZEB1 motifs in regions with increased H3K27ac in T_EX_ to regions with increased chromatin accessibility (ATAC) in T_EX_. (**d**) Heatmap showing DEGs between T_MEM_ and T_EX_ associated with regions with increased H3K27ac in T_EX_ that contain ZEB1 motifs. (**e**) Bar graph showing cell type expression of DEGs associated with ZEB1 motifs in regions with H3K27ac enriched in T_EX_ without concurrent increases in chromatin accessibility. (**f**) Genome tracks highlighting ZEB1 motifs in regions with increased H3K27ac levels in T_EX_ without changing chromatin accessibility. Differentially modified regions for H3K27ac and open chromatin are highlighted in boxes under tracks. Predicted ZEB1 binding sites are shown in red. (**g**) Number of differentially expressed TFs between T_MEM_ and T_EX_, with representative TFs indicated. (**h**) Comparison of hPTMs between T_EX_ and T_MEM_ for TFs with increased expression in T_EX_. Top six most frequent groups plotted. (**i**) Genome tracks showing RNA-seq, ATAC-seq and hPTM data for TFs with increased expression in T_EX_. Differentially modified regions for each modification are highlighted in black bars.

To further investigate the roles of ZEB1 and NUR77 at H3K27ac sites compared to accessible chromatin regions, we used motif analysis to identify predicted ZEB1 and NUR77 binding sites within differentially H3K27ac-modified regions and differentially accessible chromatin regions. Although ∼20% (38/193) of ZEB1 motifs were found in regions where both H3K27ac and chromatin accessibility were increased in T_EX_ compared to T_MEM_, ∼45% (86/193) of predicted ZEB1 binding sites had increased H3K27ac without a concurrent increase in chromatin accessibility (Fig. 2c). Of these 86 ZEB1 predicted binding sites with increased H3K27ac deposition in T_EX_, but without increased chromatin accessibility (Fig. 2c), the vast majority, ∼91%, were associated with genes increasing in expression in T_EX_ (Fig. 2d-2e). Genes potentially regulated by ZEB1 binding at sites with changing H3K27ac deposition included the TFs *Eomes* and *Tox,* and the mitotic regulator *Plk1* (Fig. 2d and 2f). In contrast, NUR77 motifs were much more prevalent in regions of increased chromatin accessibility in T_EX_ than regions with increased H3K27ac, representing 71% of predicted binding sites (Fig. S2b). We identified an enrichment in these predicted binding sites toward genes with increased expression in T_EX_ (Fig. S2c-S2d), including the genes encoding TFs *Setbp1 and Ikzf2* (HELIOS), the pro-survival factor *Bcl2* and *Tnfsf4* (OX40L) (Fig. S2c and S2e). However, ∼30% of these NUR77 motifs were located in genes highly expressed in T_MEM_. These analyses support the differential rank in Taiji PageRank analysis for ZEB1 and NUR77 and suggest a preferential role for ZEB1 in T_EX_ through binding sites marked by H3K27ac.

Assessment of TF binding motifs at H3K27ac sites revealed TFs potentially acting at putative T_EX_ enhancers. However, both the accessibility of TF binding sites and expression of these TFs themselves must be tightly regulated to orchestrate broad changes in transcriptional networks during CD8 T cell differentiation. Therefore, we next asked how the TF genes were themselves regulated by hPTMs. First, we identified TFs from the AnimalTFDB database^69^ with differential gene expression between T_MEM_ and T_EX_ cells. The majority of TFs identified in this pairwise comparison increased in expression in T_EX_, with 120 TF genes upregulated in T_EX_ versus T_MEM_ compared to only 47 TFs with increased expression in T_MEM_ (Fig. 2g). We then assessed the genes encoding these TFs for associated changes in hPTMs (Fig. 2h). For TFs that increased in expression in T_EX_ compared to T_MEM_, H3K27ac, H3K4me3 or both increased in all 6 of the hPTM patterns; however, only 2 patterns (G2 and G6) had decreased H3K27me3 and none had decreased H3K9me3 (Fig. 2h). For example, *Irf4*, *Batf* and *Ikzf2* all gained H3K27ac and H3K4me3 in T_EX_, whereas H3K27me3 was unchanged for *Batf*, but lost for *Irf4* and *Ikzf2* (Fig. 2i). Similar patterns were observed for TFs with higher expression in T_MEM_, where all of the top 6 patterns were associated with increased H3K27ac levels in T_MEM_ (Fig. S2f). Together, these analyses indicate that upregulation of TF expression is more frequently associated with an increase in the activating hPTMs rather than removal of repressive hPTMs. This pattern of regulation (gain of activation-associated modifications in T_EX_) was not unique to TFs, but was observed for all genes differentially expressed between T_EX_ and T_MEM_ (Fig. 1k). Thus, these analyses indicate that both expression of the TFs that coordinate CD8 T cell differentiation and the genes downstream of these TFs are regulated by similar patterns of hPTMs. Furthermore, these data suggest that regulating gene expression through the acquisition of activating hPTMs may be a common feature of CD8 T cell differentiation.

### Super enhancers associate with distinct transcriptional wiring of T_EX_

Super enhancers (SEs) are large clusters of enhancers with the potential to bind numerous TFs and recruit co-factors^52,70^, playing key roles in defining cell fate, controlling cell identity and/or driving disease^52^. We identified enhancers in each cell type based on the presence of H3K27ac at non-promoter regions (Fig. S3a), and ranked “stitched” enhancers using the ROSE algorithm^70,71^. Enhancers with high signal intensity were defined as super enhancers, whereas those with low signal intensity were defined as typical enhancers (TEs) (Fig. 3a and S3b). As expected, SEs were associated with higher expression of nearby genes compared to TEs (Fig. S3c). To investigate whether SEs were associated with the distinct T_EX_ and T_MEM_ cell fates, we ranked SEs within each CD8 T cell population (Fig. 3a). A number of top-ranked SEs were shared between all three CD8 T cell populations, such as *Ikzf1* (IKAROS), *Rapgef1*, *Fyn* and the transcriptional regulator *Id2*, whereas the TFs *Tbx21* (TBET)*, Zeb2* and *Runx2* were shared between T_MEM_ and T_EX_ (Fig. 3a). In contrast, the SEs near genes involved in quiescence, such as *Bach2* and *Foxp1*, ranked highly in T_N_ (Fig. 3a, left). Although several SEs were highly ranked in both T_MEM_ and T_EX_, key differences were identified. SEs more highly ranked in T_MEM_ than T_EX_ were located near genes associated with effector biology, such as *Rora*, *Klrb1b and Klrg1,* persistence-associated TFs *Bhlhe40*^68,72,73^ and *Tcf7* (TCF1), and the cytokine receptor *Il2ra* were more highly ranked in T_MEM_ than T_EX_ (Fig. 3a, middle). In contrast, SEs close to the T_EX_-associated TFs *Tox, Eomes* and *Batf*, as well as the inhibitory receptors *Pdcd1* and *Havcr2* ranked highly in T_EX_ (Fig. 3a, right). Furthermore, top-ranked SEs in T_EX_ were associated with unannotated RNAs or lncRNAs, including *2310001H17Rik* (Fig. 3a, right). Thus, H3K27ac-associated SEs likely play a role in regulating expression of key lineage-defining TFs, effector molecules and other genes driving the distinct T_N_, T_MEM_ and T_EX_ cell differentiation paths.

**Figure 3.**
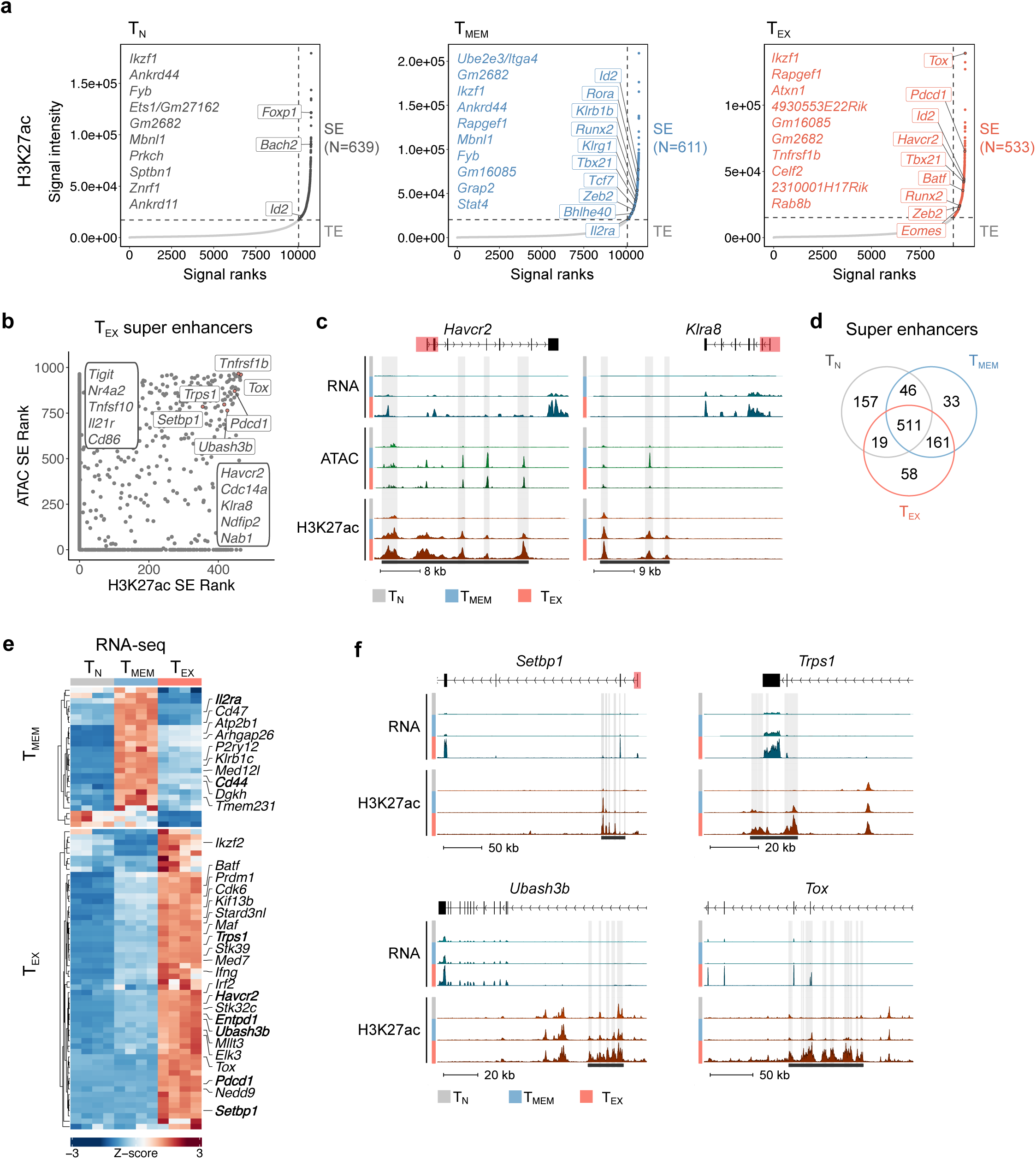
**Super enhancers drive transcriptional phenotype of T_EX_** (**a**) Distribution of H3K27ac signal across stitched enhancer regions in T_N_, T_MEM_ and T_EX_.. Top 10 ranked putative SEs plus selected SEs are highlighted. N indicates the number of putative SEs identified for each cell type. Stitched enhancers above horizontal dashed lines are associated with putative SEs; enhancers below horizontal dashed lines are typical enhancers (TE). (**b**) Comparison of SE ranks identified using H3K27ac signal (x-axis) and ATAC signal (y-axis). Selected SEs are labeled. (**c**) Genome tracks showing RNA-seq, ATAC-seq and H3K27ac data. Putative SE regions identified using H3K27ac data but not ATAC-seq data are highlighted in black boxes below tracks. Individual enhancers within SE regions are highlighted in grey. Promoter regions (±2,500bp of TSS) are indicated in red. (**d**) Venn diagram showing cell-type specificity of SEs identified by H3K27ac. (**e**) Heatmap showing gene expression of genes within 50 kb of SE identified only in T_MEM_ and T_EX_, respectively. Selected DEGs are highlighted. (**f**) Genome tracks showing RNA-seq and H3K27ac data. Putative SE regions identified using H3K27ac data are highlighted in black box. Individual enhancers within SE regions as highlighted in grey. Promoter regions (±2,500bp of TSS) are indicated in red.

Chromatin accessibility can infer SEs^52,70^ and in T_EX_, such analysis has been used to investigate the regulation of *Tox* expression^13^. Therefore, we investigated whether defining SEs by H3K27ac rather than chromatin accessibility could provide additional insights into SE regulation of the T_EX_ cell fate. We directly compared SEs defined by chromatin accessibility^5^ (Fig. S3d) to SEs identified via H3K27ac (Fig. 3a-3b). Many SEs were highly ranked using both approaches, including SEs associated with *Tox* and *Pdcd1* (Fig. 3b). However, the majority of SEs identified by chromatin accessibility were not identified using H3K27ac (Fig. S3e). For example, *Tigit*, encoding an inhibitory receptor^74^, *Nr4a2*, encoding a TF reported to promote T cell exhaustion^13,15,64^, and TNF family member *Tnfsf10*, all ranked highly for SE activity defined by chromatin accessibility but were not identified as H3K27ac-defined SEs (Fig. 3b and S3f). In contrast, *Havcr2,* encoding the inhibitory receptor TIM3, and killer cell lectin-like receptor *Klra8* ranked highly when SEs were identified by H3K27ac, but not by chromatin accessibility (Fig. 3b-3c). Furthermore, Gene Ontology (GO) analysis revealed functional divisions within SEs. Whereas GO terms for gene-associated SEs identified by chromatin accessibility were more likely to have roles in cell survival and differentiation, genes associated with SEs identified based on H3K27ac were enriched for GO terms involved in cell adhesion, division and inflammatory responses (Fig. S3g). Therefore, H3K27ac identified additional potential SE-regulated genes both with known roles in T cell exhaustion and genes that have not previously been deeply interrogated in T_EX_.

As antigen-experienced cells, T_EX_ and T_MEM_ share a core epigenetic and transcriptional network that distinguishes these cells from T_N_. However, T_EX_ and T_MEM_ also have distinct chromatin accessibility and transcriptional circuits that are cell-type specific and define these two differentiation trajectories^5,6,8,10^. More than half of all SEs we identified (52%) were shared between T_N_, T_MEM_ and T_EX_, suggesting a common role in CD8 T cell biology (Fig. 3d). Furthermore, 161 SEs were shared between T_MEM_ and T_EX_, reflecting common pathways in non-naive T cells. Moreover, only 6% of SEs were unique to T_EX_ (58/985), and even fewer were unique to T_MEM_ (3%; 33/985; Fig. 3d). Therefore, we next examined whether T cell fate-specific SEs were associated with cell type specific expression of the SE-associated gene. Indeed, T_MEM_-specific SEs were associated with high gene expression only in T_MEM_ and included *Il2ra* and *Cd44* (Fig. 3e). In contrast, genes with SEs specific to T_EX_ were highly expressed only in T_EX_, including the SE-associated genes *Tox* and the inhibitory receptors *Entpd1*, *Pdcd1* and *Havcr2* (Fig. 3e-3f). This analysis also revealed T_EX_-enriched SE-associated genes that have not been extensively studied in T_EX_, including *Setbp1*^13^, *Trps1* and *Ubash3b* (Fig. 3e-3f). Together, these data add further support to the hypothesis that SEs play key roles driving both the shared and distinct chromatin regulatory and transcriptional circuitry of T_EX_ versus T_MEM_ cells and identify previously understudied H3K27ac-enriched SEs associated with genes in T_EX_ that warrant further investigation.

### Chromatin state analysis reveals state-specific transitions from T_N_ to T_MEM_ or T_EX_

To study how genome-wide chromatin states change during CD8 T cell differentiation, we applied ChromHMM, an algorithm that uses multiple hPTMs to segment the genome into distinct states^75^. We used ATAC-seq, H3K27ac, H3K4me3, H3K27me3, and H3K9me3 data to identify the four major promoter states; active (I), poised (II), repressive (III) and repetitive/heterochromatin (IV) promoters. Active promoters (I) were defined by open chromatin and by the active marks H3K27ac and H3K4me3 (Fig. S4a-S4b). As expected, the vast majority of (∼95%) of genes with active promoters in one CD8 T cell population were highly expressed in that cell type (Fig. S4c). Pathway analysis of genes with active promoters in T_N_ revealed that these promoters were associated with core cellular function pathways, including DNA repair, chromatin segregation and translation (Fig. S4d), suggesting that, in T_N_, genes involved in basic cellular functions are regulated by promoters with active marks. In contrast, repressed (III) and repetitive/heterochromatin (IV) promoters were defined by deposition of the repressive marks H3K27me3 and/or H3K9me3 (Fig. S4a-S4b). Accordingly, the vast majority of genes with repressed or repetitive/heterochromatin promoters were not expressed in the cell type with those repressed or repetitive/heterochromatin features (Fig. S4c). In T_N_, genes with repressed promoters were enriched for more specialized pathways with limited roles in CD8 T cells, such as muscle contraction, sensory perception of pain and response to pheromone (Fig. S4d).

The poised promoter state was first described in embryonic stem cells (ESCs) as nucleosomes bearing both H3K4me3 and H3K27me3^76–78^. The genes associated with these dual modified promoters were not expressed in ESCs, but instead were turned on as cells acquired identity and lineage commitment in the developing embryo^79,80^. This poised state has also been described in “multipotent” T_N_^81^. To investigate how genes with poised promoters are associated with T_MEM_ and T_EX_ cell fates, we first identified genes with poised (state II) promoters in T_N_ (Fig. S4a-S4b). As expected, in T_N_ the majority (∼57.7%) of genes with promoters bearing both H3K4me3 and H3K27me3 were not highly expressed (Fig. S4c), supporting the assignments of these promoters as poised. Within the 4,945 poised T_N_ promoters, we identified promoters that shifted to an activated state only in T_EX_ or only in T_MEM_ (Fig. 4a, indicated by star and triangle respectively). We then focused on promoters that were associated with increased gene expression in each cell type (Fig. 4b). For example, the Src family kinase *Yes1* and the IL-2 receptor alpha (*Il2ra*) showed this pattern of activation only in T_MEM_ (Fig. 4b-4c). Furthermore, promoters for genes posited to have roles in CD8 T cell persistence and tissue residency, including *Prss12*^82^, *Nt5e* (CD73^83^), and *Ier3* (IEX-1^84^) transitioned from poised-to-active state only in T_MEM_ (Fig. 4b). Gene ontology (GO) analysis revealed functions in metabolism and signaling, including cytokine signaling (Fig. S4e). Together these analyses suggest that genes poised in T_N_ that become activated in T_MEM_ are involved in key aspects of T_MEM_ biology.

**Figure 4.**
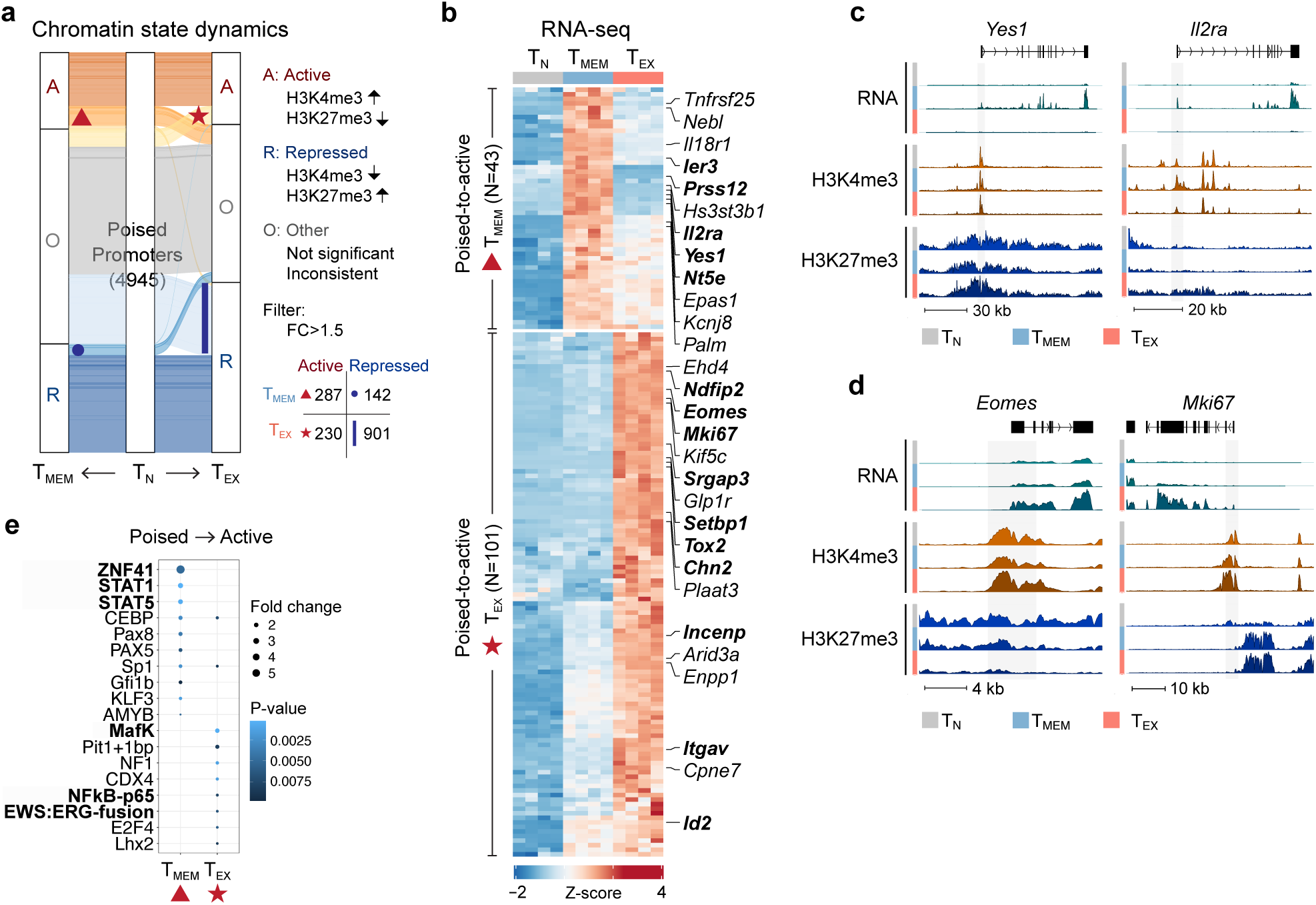
**Key T_MEM_ and T_EX_ genes are poised in T_N_ and activated upon differentiation** (**a**) Alluvial plot showing how hPTMs at promoters poised in T_N_ change in T_MEM_ and T_EX_. Active promoters were defined as either significantly gaining H3K4me3 or losing H3K27me3 or both, repressed promoters were defined as either significantly losing H3K4me3 or gaining H3K27me3 or both. Statistical cutoff of FC > 1.5. Number represents total number of T_N_ poised protomers; inset table shows number of poised-to-active and poised-to-repressed promoters in T_MEM_ and T_EX_. (**b**) RNA expression heatmap for DEGs with poised promoters which were activated in either T_MEM_ (triangle) or T_EX_ (star). Selected DEGs are highlighted in bold. (**c-d**) Genome tracks showing poised promoters in T_N_ that switched to active **(c)** only in T_MEM_ or **(d)** only in T_EX_. Promoters were excluded from analysis and are highlighted in grey bars. (**e**) Bubble plot showing changes in predicted TF binding site accessibility for poised-to-active T_MEM_ or T_EX_ genes, with select TFs highlighted.

In contrast, genes with promoters that shifted from T_N_ poised to active in T_EX_ included the TFs *Eomes*, which drives terminal differentiation of T_EX_^19,61^ and *Tox2*, which has a reported role in T_EX_^15^, as well as other T_EX_-associated genes including *ItgaV* (CD51)^61^, *Srgap3*^85^, and *Ndfip2*^85^ (Fig. 4b and 4d). Several cell cycle-associated genes also showed this pattern of regulation, including *Mki67* (KI67), *Incenp* and *Chn2* (Fig. 4b and 4d) consistent with the more extensive cell division history of T_EX_ including ongoing cell cycle^10,19^. Of note, the promoter of *Id2* was poised in T_N_, and shifted to an activated state only in T_EX_, despite RNA expression in both T_MEM_ and T_EX_ (Fig 4b^86–88)^. Additional analysis revealed that, although H3K27me3 was lost at the *Id2* promoter in both T_MEM_ and T_EX_, T_EX_ retained activating H3K4me3 and this modification decreased in T_MEM_ compared to T_EX_ (Fig. S4f). These analyses highlight the complexity of gene expression regulation by a suite of hPTMs at and around the promoter. Finally, several genes that have not previously been well studied in T_EX_ were also identified, including *Setbp1* (Fig. 4b), which was also associated with a SE in T_EX_ (Fig. 3e-3f). GO analysis identified pathways linked to development, differentiation, and proliferation (Fig. S4e). Together, these data suggest that genes with poised promoters in T_N_ are key genes involved in the divergent differentiation trajectories of T_MEM_ and T_EX_, including lineage-driving TFs and genes associated with T_EX_ function (e.g. continued proliferation and survival during high antigen stress).

We next investigated TFs associated with expression of genes with poised-to-active promoters and might therefore drive the distinct T_EX_ and T_MEM_ cell fates. We performed TF motif analysis on these poised-to-active promoters (Fig. 4a-4b). Motifs for ZNF41, a zinc finger family TF, were enriched in T_MEM_ poised-to-active promoters, as were both STAT1 and STAT5 motifs (Fig. 4e). Indeed, STAT1 is required for CD8 T cell clonal expansion and memory formation^89^, and STAT5 has a key role in early effector and memory-precursor CD8 T cell differentiation^90^. In contrast, predicted MafK binding sites were enriched in poised-to-active T_EX_ promoters (Fig. 4e). MafK forms heterodimers with BACH2 to help direct the repressive activity of BACH2 to specific genes^91^. BACH2 is a key transcriptional coordinator involved in T cell quiescence in T_N_, T_MEM_ including stem cell memory cells, but also in stem cell-like progenitor T_EX_^92,93^ suggesting a potential T_EX_-associated BACH2-MafK regulatory module. Furthermore, Fli1, an ETS family TF, dampens effector CD8 T cell transcriptional networks^94^. The EWS:ERG fusion motif, which is also predicted to be bound by Fli1, was enriched in poised-to-active promoters in T_EX_ (Fig. 4e), supporting a potential role for Fli1 or other ETS family members in restraining effector biology in T_EX_. Finally, NFkB-p65 motifs were enriched in T_EX_ poised-to-active promoters (Fig. 4e). NFkB signaling has broad roles in T cells, regulating initial TCR-mediated T cell activation, proliferation and effector function as well as T_MEM_ survival^95,96^. Moreover, NFkB transcriptional circuitry is augmented after treatment with immunotherapies targeting inhibitory receptors, such as PD-1 blockade, or costimulation, such as CD137 (41BB) agonism^5,97^. The enrichment for NFkB-p65 motifs at T_EX_ poised-to-active promoters suggests that reengaged NFkB circuitry in immunotherapy-reinvigorated T_EX_ may be driving expression of previously poised genes, with potential implications for improving therapies. Together, these data suggest that a subset of the distinct T_EX_ and T_MEM_ transcriptional programs consist of genes with poised promoters in T_N_ that then acquire either active or repressed chromatin states during differentiation.

We next investigated how chromatin modifications could reinforce distinct T_MEM_ and T_EX_ cell states through repression of alternative fates. We identified promoters that were poised in T_N_ and then became repressed in T_MEM_ or T_EX_ either through gain of H3K27me3, loss of H3K4me3 or both potentially to silence genes of alternative cell fates (Fig. 4a). In total, 901 promoters switched from a poised to repressed state in T_EX_ compared to only 142 promoters for T_MEM_ (Fig. 4a, blue line for T_EX_ compared to blue circle for T_MEM_). Genes with promoters that switched from poised-to-repressed as T_N_ differentiated into T_MEM_ on average lost the activating mark H3K4me3 and also maintained the repressive mark H3K27me3 (Fig. S4g). For example, several genes associated with T cell differentiation including the TFs *Tox2, Eomes* and *Ikzf2* (HELIOS), the exhaustion-associated ectonuclease *Entpd1* (CD39), costimulatory molecule *Tnfsf4* (OX40L) and glycoprotein *Itm2a* all had higher H3K27me3 and lower H3K4me3 at the promoters for these genes in T_MEM_ compared to T_N_ and T_EX_ (Fig. S4h). This poised-to-repressed promoter state was associated with lower RNA expression of these genes in T_MEM_ than in T_EX_. (Fig. S4i), indicating that genes highly expressed in T_EX_ and poised in T_N_ are actively repressed in T_MEM_. In contrast, genes that became repressed during T_EX_ differentiation gained repressive H3K27me3 at the promoter, however there was variable loss of H3K4me3 at these promoters (Fig. S4j). For example, in T_EX_, whereas *Ier3* (IEX1)*, Tnfrsf25* (DR3)*, Hdac10* and *Mapk12* gained H3K27me3 at the promoter, only *Mapk12* lost H3K4me3 (Fig. S4h, S4j-S4k). Furthermore, a subset of genes with promoters that switched from poised-to-repressed in T_EX_ switched from poised-to-active in T_MEM_ (e.g. *Ier3*, *Il2ra*, *Tnfrsf25*) and vice versa (e.g. *Tox2*) (Fig. 4b and S4h). Together, these data suggest that active repression of genes poised in T_N_ may help enforce the distinct T_MEM_ and T_EX_ states and potentially limit conversion between these two populations, with gain of H3K27me3 predominantly driving this repression in T_EX_.

### Atypical H3K9me3 narrow peaks are enriched for CTCF motifs and occur at distinct repeat classes

H3K9me3 is typically associated with constitutive heterochromatin and is classically involved in silencing gene expression, playing a critical role in cell differentiation^98,99^. This hPTM classically exhibits broad peaks across the genome^98^, however analysis of H3K9me3 peak width in CD8 T cells revealed a wide range in peak size, from <5 kb to >100 kb, with most peaks less than 10 kb (Fig. S5a)^100,101^. Therefore, we examined how H3K9me3 localization and deposition changed during CD8 T cell differentiation. Regions with higher H3K9me3 in T_MEM_ compared to T_EX_ (T_MEM_-enriched peaks; n=1565) were broad, covering ∼40.1kb bases on average (Fig. 5a, left; Fig 5b). In contrast, regions with higher H3K9me3 in T_EX_ compared to T_MEM_ (T_EX_-enriched peaks; n=1279) were narrower, averaging only ∼15.7kb bases (Fig. 5a, right; Fig. 5c). Directly comparing the distribution of peak sizes between T_EX_-enriched and T_MEM_-enriched H3K9me3 peaks showed that T_EX_-enriched peaks were substantially narrower than T_MEM_-enriched peaks (Fig. 5d). Moreover, only 10.2% (n=130) of T_EX_-enriched peaks were broad (>=15 kb), compared to 42.9% (n=672) of T_MEM_-enriched peaks (Fig. 5e). Analysis of the number of base pairs in the genome covered by H3K9me3 revealed that while the majority of the genome is covered by broad peaks, over 25% of the T_EX_-enriched base pairs are in narrow peaks (Fig. S5b). The majority of narrow T_EX_-enriched H3K9me3 peaks were found in intergenic and intronic regions, with only a small proportion (∼3%) located in promoter regions (Fig. S5c). These results suggest that H3K9me3 deposition patterns at T_MEM_ or T_EX_-enriched peaks have distinct characteristics, with H3K9me3 enriched in narrow peaks in T_EX_ and broad peaks in T_MEM_.

**Figure 5.**
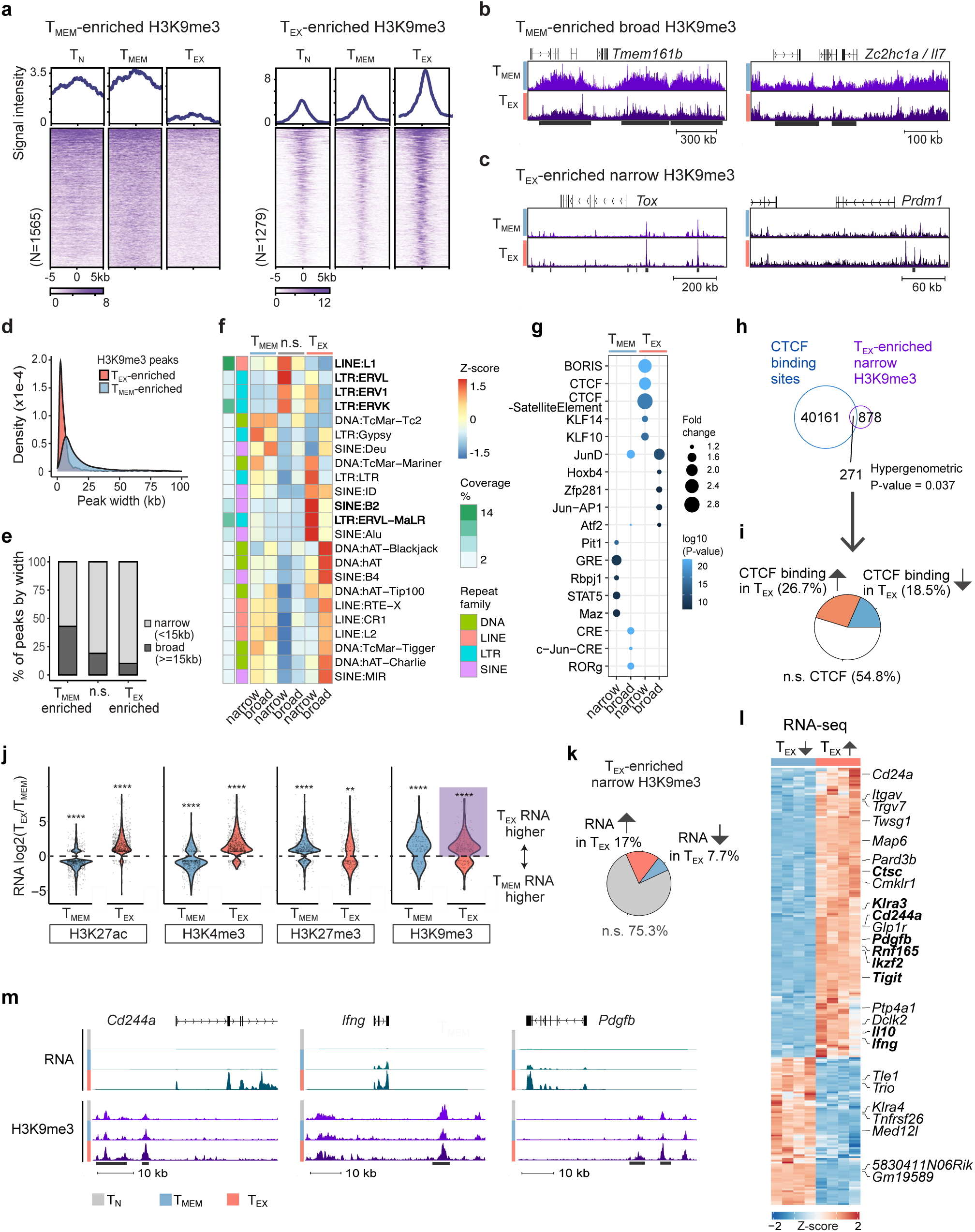
**T_EX_-enriched atypical H3K9me3 peaks cover CTCF sites and are associated with gene expression** (**a**) Signal intensity heatmap of T_MEM_-enriched and T_EX_-enriched H3K9me3 regions. (**b**) Genome track showing broad H3K9me3 regions enriched in T_MEM_. (**c**) Genome track showing narrow H3K9me3 regions enriched in T_EX_. (**d**) Peak size distribution of T_MEM_- and T_EX_-enriched H3K9me3 regions. (**e**) Percentage of H3K9me3 peaks that are broad versus narrow. Peaks >=15 kb are defined as broad, peaks <15 kb are defined as narrow. N.s. = non-significantly different between T_MEM_ and T_EX_. (**f**) Heatmap of Z-scored repeat element class coverage of T_MEM_-enriched, non-significant (n.s.) and T_EX_-enriched H3K9me3 narrow and broad peaks. (**g**) Predicted TF binding motifs in T_MEM_- and T_EX_-enriched narrow and broad peaks compared to n.s. H3K9me3 peaks. Top 5 motifs shown per comparison. (**h**) Venn diagram showing overlap between locations of CTCF binding sites and T_EX_- enriched narrow H3K9me3 peaks. P value represents a hypergeometric test for the comparison. (**i**) Pie chart showing change in CTCF binding at sites within T_EX_-enriched narrow H3K9me3 peaks. (**j**) Violin plot of log2 fold change in RNA expression between T_MEM_ and T_EX_ near differentially modified regions for each hPTM (H3K27ac, H3K4me3, H3K27me3 and H3K9me3). (**k**) Pie chart showing changes in RNA expression between T_MEM_ and T_EX_ of genes near T_EX_-enriched narrow H3K9me3 peaks. (**l**) Heatmap of RNA expression for genes from Fig. 5k. (**m**) Genome track showing DEGs near to narrow T_EX_-enriched H3K9me3 regions.

A major function for H3K9me3 is to repress repetitive genomic elements, maintaining genome integrity^102^. Therefore, we examined whether the atypical narrow H3K9me3 peaks enriched in T_EX_ also were associated with repetitive elements like broad H3K9me3 peaks. We first assessed repeat coverage of narrow, broad and H3K9me3 peaks that were not significantly different (n.s.) between T_EX_ and T_MEM_. Analysis of T_EX_-enriched H3K9me3 peaks shows that 53% of narrow peaks covered repeats, compared to around 34% of broad peaks (Fig. S5d). This analysis suggests that broad and narrow H3K9me3 peaks may function to repress repetitive genomic elements, but that narrow H3K9me3 peaks could be additionally specialized to this role.

We next investigated whether specific families of repetitive elements were enriched underneath the T_EX_-enriched narrow H3K9me3 peaks. T_EX_-enriched narrow H3K9me3 peaks had reduced coverage of LINE elements (27.7%) (Fig. S5e, top) compared to n.s. narrow peaks (60.8%) and T_MEM_-enriched narrow peaks (52%), as did T_EX_-enriched broad peaks (Fig. S5e, bottom). However, T_EX_-enriched narrow peaks showed an increase in LTR element coverage compared to other H3K9me3 peak sets (Fig. S5e). To further investigate this finding, we assessed repeat element subclasses. Narrow H3K9me3 peaks that were not enriched in T_MEM_ or T_EX_ were strongly associated with LINE:L1 and LTR:ERV subclasses compared to n.s. broad H3K9me3 peaks (Fig. 5f). T_MEM_-enriched broad and narrow peaks had similar patterns of repeat element coverage, wheras T_EX_-enriched broad and narrow peaks were associated with distinct repeat element subclasses (Fig. 5f). For example, T_EX_-enriched narrow H3K9me3 peaks showed a unique enrichment in the LTR subclass ERVL-MaLR and the retrotransposon SINE B2 (Fig. 5f), suggesting a distinct regulatory role for these elements in T_EX_ cells.

Repeat elements have been co-opted to serve critical roles in chromatin organization, enhancer function, and gene regulation^103,104^. SINE B2 is one example of repeat elements serving functional roles. These elements are rodent-specific retrotransposons that contain binding sites for CTCF^105^, a zinc finger protein that acts as a transcriptional activator, repressor, and genome organization^106^. H3K9me3 at SINE B2 repetitive elements can regulate CTCF binding at these sites^107^. Therefore, we performed an unbiased motif analysis of T_MEM_- and T_EX_-enriched H3K9me3 peaks. Indeed, the top most enriched motifs under T_EX_ narrow peaks were CTCF and the CTCF related factor (CTCFL or BORIS) (Fig. 5g). To validate whether CTCF bound directly at T_EX_-enriched H3K9me3 narrow peaks, we performed CUT&RUN for CTCF in T_N_, T_MEM_, and T_EX_. Nearly a quarter (24%, n=271) of T_EX_-enriched H3K9me3 narrow peaks were bound by CTCF (Fig. 5h, p = 0.037). Of these, 45% had differential CTCF binding: 26.7% showed increased CTCF binding in T_EX_ compared to T_MEM_, while 18.5% had reduced CTCF binding in T_EX_ (Fig. 5i). For example, CTCF binding increased in H3K9me3 peaks associated with *Ctsc* (CAPTHESIN C), a peptidase that coordinates activation of serine proteases including granzymes, whereas CTCF binding decreased at a H3K9me3 peak close to *Klra3*, an NK cell receptor expressed in a cytotoxic subset of T_EX_ (Fig. S5f). Together, these findings suggest that narrow H3K9me3 peaks may have distinct associations with repetitive elements in different CD8 T cell subtypes, and that altered H3K9me3 deposition in T_EX_ may impact CTCF binding at specific sites in the genome, with potential implications for cell type-specific genome organization.

To investigate whether these T_EX_-enriched narrow H3K9me3 peaks were associated with chromatin accessibility, we examined the overlap between these peaks and T_EX_-accessible chromatin regions. Approximately 25% (287/1149) of T_EX_-enriched narrow H3K9me3 peaks were located in regions of open chromatin in T_EX_ (Fig. S5g), for example near *Tox* (Fig. S5i). Genes with narrow H3K9me3 peaks associated with open chromatin, were significantly enriched in cytokine-mediated signaling and leukocyte cell-cell adhesion pathways (Fig. S5h). These observations suggests that the association of a subset of narrow H3K9me3 peaks with open chromatin may contribute to the role for this hPTM to regulate expression of key cell-type associated genes.

### T_EX_-enriched H3K9me3 peaks correlate with gene activation

H3K9me3 is typically associated with repression of gene expression^108^. However, a subset of genes with increased expression in T_EX_ compared to T_MEM_ had higher H3K9me3 deposition in T_EX_ (Fig. 1k, G5 and G6). Thus, we next investigated the association between H3K9me3 and gene expression in T_EX_. Regions with differentially enriched hPTMs between T_MEM_ or T_EX_ were identified, then filtered for nearby genes that were differentially expressed between T_MEM_ and T_EX_ (Fig. 5j). As expected, genes near regions with increased H3K27ac or H3K4me3 in one cell type had overall higher expression in that cell type (Fig. 5j), consistent with gene-activating functions of H3K27ac and H3K4me3. In contrast, genes near regions of increased H3K27me3 or H3K9me3 in T_MEM_ had lower expression in T_MEM_ than T_EX_ (Fig. 5j), consistent with the repressive roles of H3K27me3 and H3K9me3. However, although increased H3K27me3 was negatively associated with gene expression in T_EX_, H3K9me3 was positively associated with gene expression, with nearby genes more highly expressed in T_EX_ versus T_MEM_ (Fig. 5j, purple box). Given the observations above that increased H3K9me3 deposition in T_EX_ was predominantly localized to narrow peaks (Fig 5a and 5e; 89.8% of peaks), we tested if the link between H3K9me3 and increased gene expression was associated with narrow H3K9me3 peaks, or a general feature of H3K9me3 in T_EX_. Both narrow and broad H3K9me3 peaks showed the same pattern of increased deposition of H3K9me3 near genes with higher expression in T_EX_ (Fig. S5j).

To assess the breadth of impact of these H3K9me3 peaks on regulating the transcriptional programs of T_EX_, we next asked what proportion of T_EX_-enriched peaks were associated with changes in gene expression (Fig. 5k). Given that the vast majority of H3K9me3 peaks in T_EX_ were narrow, we focused the subsequent analyses on these atypically narrow peaks. The majority of genes close to T_EX_-enriched H3K9me3 narrow peaks did not change expression (n.s., Fig. 5k), suggesting that not all H3K9me3 has gene regulatory functions. However, 17% of H3K9me3 T_EX_-enriched narrow peaks were associated with gene upregulation in T_EX_ cells, whereas 7.7% were associated with decreased expression (Fig. 5k), such that narrow H3K9me3 T_EX_-enriched peaks were twice as likely to be associated with gene upregulation compared to downregulation. Of note, T_MEM_-enriched narrow H3K9me3 peaks did not show this positive association with gene expression (Fig. S5k). Furthermore, 12.3% of genes upregulated in T_EX_ compared to T_MEM_ had an increase in H3K9me3 deposition in a narrow peak within 50 kb (Fig. S5l). This fraction increased to 34% of genes when assessing H3K9me3 deposition within 250 kb of the gene (Fig. S5l), suggesting that a subset of T_EX_ upregulated genes were potentially regulated at least in part by atypical “activating” H3K9me3.

Finally, we performed a GO analysis to examine differences in the biological processes associated with H3K9me3 deposition in T_EX_. For example, genes nearby T_EX_-enriched narrow peaks with decreased expression in T_EX_, indicative of a repressive function for H3K9me3, were enriched for negative regulation of lymphocyte activation (Fig. S5m). In contrast, genes with increased expression close to regions with increased H3K9me3 deposition in T_EX_, i.e. “activating” H3K9me3, were associated with cell cycle, cell death, response to cytokine stimulus and leukocyte activation (Fig. S5m). Thus, genes potentially regulated by both canonical repressive and non-canonical “activating” H3K9me3 play broad roles in T_EX_ biology. Indeed, upregulated genes in T_EX_ located near T_EX_-enriched narrow H3K9me3 peaks included the key T cell exhaustion TFs *Tox, Prdm1* (BLIMP-1) and *Ikzf2*, inhibitory receptors *Cd244a* (2B4) and *Tigit*, and functional molecules *Ifng*, *Il10* and *Pdgfb* (Fig. 5c, 5l and 5m). Together, these results suggest that the H3K9me3 modification may have distinct characteristics in T_EX_, with increased deposition localized to non-conventional narrow peaks, a subset of which are located near key genes that increase in expression in T_EX_, including *Tox*.

## DISCUSSION

Here we profiled the hPTMs and chromatin epigenetic landscape of CD8 T cells as they differentiate from T_N_ into two functionally different cell fates, T_MEM_ and T_EX_. T_N_, T_MEM_ and T_EX_ cells had distinct epigenetic profiles across both activating (H3K27ac and H3K4me3) and repressive (H3K27me3 and H3K9me3) hPTMs, with the majority of hPTM changes occurring as T_N_ were activated and differentiated into T_MEM_ or T_EX_. The unique transcriptional networks of T_MEM_ and T_EX_ were co-regulated by combinations of hPTMs, with gain of activating hPTMs playing a dominant role compared to loss of repressive modifications. Differentiation from T_N_ into T_MEM_ or T_EX_ resulted in both activation and repression of discrete subsets of genes that were poised in T_N_, suggesting that hPTMs play a role in both upregulating T_MEM_ versus T_EX_ transcriptional networks, and repressing transcription of genes associated with the opposing cell fate. Whereas increased deposition of H3K9me3 in T_MEM_ was associated with decreased gene expression in T_MEM_, a subset of genes with increased expression in T_EX_, including the TF *Tox*, had increased H3K9me3 in nearby non-canonical narrow peaks. Thus, our analyses reveal the complexity of hPTMs in guiding alternative CD8 T cell fates, with potentially atypical roles in T_EX_.

T_EX_ have a unique open chromatin accessibility and transcriptional landscape compared to T_MEM_. However, precisely how this cell-fate specific chromatin accessibility may mediate cell-fate associated gene expression remains poorly understood. Analysis of individual hPTMs revealed that in both T_MEM_ and T_EX_ gain of activating modifications in one cell type was strongly associated with increased gene expression in this cell type. In contrast, loss of repressive modifications was only loosely correlated with increased gene expression in this cell type. Supporting this observation, we found that activating modifications were frequently gained in the most common combinatorial patterns of hPTMs associated with cell-type specific increases in gene expression, but that often these changes in H3K27ac and H3K4me3 did not co-occur with loss of H3K27me3 and/or H3K9me3. Thus, these analyses suggest that active modifications are key components for gene activation in CD8 T cells, whereas repressive hPTMs may serve a fine-tuning role, selectively regulating a subset of potentially cell-fate related genes.

To further investigate how combinations of activating and repressive marks regulate T_MEM_ and T_EX_ gene expression, we focused on poised chromatin states (with both H3K4me3 and H3K27me3) and the dynamics of these states as CD8 T cells differentiate. Genes in a poised state displayed high cellular plasticity, enabling them to quickly respond to antigen stimulation and facilitate rapid cell differentiation^109–111^. Thus, poised chromatin states are linked to genes driving cellular identity^110^. Since T_MEM_ and T_EX_ share a common progenitor (T_N_), we investigated how genes in a poised state in T_N_ might shape the distinct transcriptional networks of these populations. Discrete subsets of genes poised in T_N_ shifted to an active state in either T_MEM_ or T_EX_. In T_MEM_, this subset included genes such as *Ier3* and *Il2ra,* and in T_EX_ included multiple TFs such as *Eomes*, *Setbp1*, *Tox2* and *Id2* in addition to genes such as *Ki67*. Furthermore, genes that shifted from poised-to-active promoter states only in T_MEM_ or only in T_EX_ were regulated by distinct families of TFs, with STAT family binding motifs enriched in T_MEM_ poised-to-active genes compared to MafK and NFkB-p65 in T_EX_. Thus, the distinct functions of T_MEM_ and T_EX_ were regulated by discrete sets of TFs that coordinated upregulation of key genes primed for activation in quiescent T_N_. Whether this poised state is imprinted during thymic development when many T cell genes are “tested” or reflects even earlier developmental poising will be interesting to examine in the future. Furthermore, T_MEM_ and T_EX_ are distinct endpoints in complex differentiation trajectories originating from a common T_N_ precursor. It will be interesting to investigate the dynamics of how promoters poised in T_N_ are activated/repressed throughout the full trajectory of T_MEM_ and T_EX_ differentiation, for example as early T_MEM_ precursors differentiate into T_MEM_. T_EX_ are epigenetically inflexible and do not convert to T_EFF_ or T_MEM_ cell states. Analysis of genes that were poised in T_N_ but shifted to a repressed state in T_MEM_ or T_EX_ highlighted the role of active repression in forming and maintaining these distinct CD8 T cell populations. For example, expression of the inhibitory receptor *Entpd1* was repressed in T_MEM_, whereas *Il2ra* was repressed in T_EX_. Together, these data suggest that, although gain of activating hPTMs plays a dominant role in the upregulation of discrete TF regulatory networks in T_MEM_ and T_EX_, repressive hPTMs also coordinate the repression of opposing CD8 T cell fates. These findings highlight the multidimensional roles of hPTMs in T_EX_, and emphasize the importance of both activating and repressive modifications. Understanding the combinations of these hPTMs provides a comprehensive view of the regulatory landscape, and reveals the complex patterns regulating gene expression in T_EX_.

The top three “patterns” of hPTM changes associated with gene upregulation were the same between T_MEM_ and T_EX_, indicating that the broad associations of these hPTMs and gene expression are comparable between CD8 T cell states. However, in T_EX_, a subset of upregulated genes was associated with gain of both H3K27ac and H3K4me3, but also a gain of H3K9me3, typically a repressive modification. These genes included key T_EX_ TFs, such as *Tox* and *Ikzf2*, effector genes including *Ifng,* and the gene including the inhibitory receptor *Cd244a* (2B4). This finding was in contrast to the widely understood role of H3K9me3 and its association with gene repression, especially in embryonic stem cells^108,112^, suggesting that this modification may have a distinct function in T_EX_. Indeed, we found that classically broad H3K9me3 was, as expected, associated with decreased gene expression in T_MEM_, indicating that H3K9me3 performs this typical role in T_MEM_. However, in T_EX_, H3K9me3 was predominantly gained in atypically narrow peaks and, furthermore, these narrow H3K9me3 peaks were associated with gene activation. Narrow H3K9me3 peaks were enriched for CTCF motifs and the repetitive elements SINE:B2 and LTR:ERVL-MaLR, suggesting that CTCF binding could be regulated by H3K9me3 deposition in T_EX_. Specifically, increased H3K9me3 at SINE B2 sites in T_EX_ may influence CTCF binding patterns and therefore genome organization during CD8 T cell exhaustion, potentially contributing to increased gene expression at these locations. These data provoke the hypothesis that gene activation in T_EX_ is, at least in part, mediated by atypical H3K9me3 deposition that influences CTCF binding to alter higher order chromatin structure, which may in turn regulate gene expression. This result highlights an unusual role of H3K9me3 in gene regulation in T_EX_. Understanding whether and how this role impacts chromatin organization, and what the relationship is between these hPTM patterns and CTCF binding, for example, in T_EX_ biology will be of interest for future studies.

Multiple recent studies have used ATAC-seq to examine the unique chromatin accessibility landscape of T_EX_ and to investigate how this landscape impacts T_EX_ biology^5,6,8,10,7,9,113,114^. However, chromatin accessibility is only one feature of a dynamic epigenetic landscape and analysis of hPTMs may provide additional insights into CD8 T cell subset differentiation and T_EX_ biology. Using H3K27ac deposition to identify cell-state associated SEs uncovered additional SEs compared to examination of chromatin accessibility alone. For example, we discovered a potential role for SEs in regulating expression of *Havcr2* (TIM3) and *Klra8* using H3K27ac, whereas ATAC-seq did not identify these SEs. In addition, analysis of predicted TF activity at H3K27ac sites provided further insight into TF function at enhancers and SEs. For example, the TF NUR77 was highly ranked in T_EX_ compared to T_MEM_ only when activity was predicted using open chromatin data, but not when H3K27ac-decorated regions were analyzed. This observation suggests that NUR77 may be predominantly acting in T_EX_ at genomic locations not associated with enhancer or SE activities. In contrast, the TF ZEB1 was predicted to have high importance in T_EX_ compared to T_MEM_ at H3K27ac-associated enhancers and SEs, but not across open chromatin regions in general. Thus, understanding the additional layer of regulatory networks modulated by hPTMs on top of chromatin accessibility provides further insight into how the distinct T_MEM_ and T_EX_ transcriptional programs are established. Further work is required to investigate how TFs such as ZEB1 function at SEs.

In this study, we interrogated the epigenetic landscape of two distinct CD8 T cell fates, T_MEM_ and T_EX_, and their common precursor T_N_. Despite these two populations representing endpoints of an antigen-driven differentiation hierarchy, T_MEM_ and T_EX_ themselves are heterogeneous and contain further proliferative and functional hierarchies, including subsets that function as stem cell-like reservoirs for more terminally differentiated, effector-like populations^19–25,18,10^. It is likely that observations made for bulk T_EX_ and T_MEM_ populations reflect an average hPTM landscape and average gene expression across distinct T_MEM_ and T_EX_ subsets. For example, it is likely that additional hPTM associations with transcriptional circuits will be identified in subsets of T_EX_ such as the key progenitor T_EX_ and the downstream T_EX_ intermediate and terminal populations. This subset-specific variation in hPTMs could contribute to functional differences between T_EX_ subsets, potentially shaping the diversity of T cell responses. Similar heterogeneity exists in T_MEM_ as well. Thus, future research should explore how hPTMs mediate control of gene expression within these additional T_MEM_ and T_EX_ subsets, including at earlier timepoints in the differentiation trajectories of these subsets. Our analysis of hPTMs uncovered a potential interplay between hPTM deposition and higher-order chromatin structure in regulating gene expression in T_EX_. How the deposition of these hPTMs, including atypical H3K9me3 in T_EX_, is orchestrated remains unknown, including the role of T_MEM_ and T_EX_ TF networks in recruiting epigenetic enzymes to sites of both histone modification and chromatin remodeling. It will be interesting in the future to investigate how TF networks, hPTMs, and three-dimensional genome structure coordinate establishment and maintenance of the distinct transcriptional landscape of T_MEM_ and T_EX_ and their subsets. Thus, understanding how the T_MEM_ and T_EX_ differentiation hierarchies are epigenetically regulated will provide key insight into the epigenetic scar of exhaustion, fate-flexibility, and could be used to inform effective clinical therapies.

## MATERIALS AND METHODS

### Mice

Animals were housed in a specific pathogen-free facility at the University of Pennsylvania at ∼20°C with 55% humidity and a dark-light cycle of 12hr-12hr. Animals were provided with ad libitum access to food and water throughout the duration of the experiment. All experiments and breeding were approved by the Institutional Animal Care and Use Committee guidelines for the University of Pennsylvania. All procedures were performed in accordance with Institutional Animal Care and Use Committee Protocol 803071. Transgenic mice expressing a TCR specific for the LCMV peptide D_b_GP_33-41_ (P14 donor mice) were bred in-house at the University of Pennsylvania on a C57BL/6 background purchased from Charles River. Donor mice were used at ∼8 weeks of age. Recipient C57BL/6 mice were purchased from Charles River and used at 6-8 weeks of age. Recipient mice were sex matched with donor mice. Euthanasia was performed using CO_2_ inhalation in a CO_2_ unit as recommended by the Panel on Euthanasia of the American Veterinary Medical Association and the University of Pennsylvania.

### Infections

LCMV Armstrong (Arm) and LCMV clone 13 (Cl13) were grown in house and titrated as previously described^5^. Recipient mice were either infected intraperitoneally (i.p) with 2 x 10^5 PFU LCMV Armstrong to model an acute infection or intravenously (i.v.) with 4 x 10^6 PFU LCMV Cl13 to establish a chronic infection.

### Adoptive cell transfer

PBMCs were isolated from the peripheral blood of naive P14 donor mice using gradient centrifugation (Histopaque-1083). 1,000 naive P14 cells were adoptively transferred i.v. into sex-matched recipient mice. P14 cells were isolated from donor mice of a distinct congenic background than recipient mice to enable donor P14 cells to be distinguished from recipient CD8 T cells. Recipient mice were infected with LCMV Arm or LCMV Cl13 one day following adoptive cell transfer.

### Cell sorting

Spleens were collected at d30 of LCMV Arm and d32 of LCMV Cl13 infection. Donor P14 mice or littermates were used for the naive condition. Single cell suspensions were prepared by mechanical disruption of spleens through a 70 µm cell strainer. Red blood cells were lysed in ACK buffer (3min, RT) and CD8 T cells isolated using EasySep CD8 T cell negative selection kit (Stem Cell, Cat# 19853) following manufacturer’s instructions. CD8 T cells were washed in FACS buffer (2% FCS in PBS) and surface stained with an antibody cocktail in FACS buffer for 30 min at 4°C. Donor CD8+ P14 cells were sorted on a BD FACS Aria II using congenic markers for identification. Samples were sorted to >95% purity.

## CUT&RUN

CUT&RUN was performed as previously described with slight modifications^94,115^. 10,000 sorted cells were washed twice (600 g x 5 min) with 1 ml of cold wash buffer (20 mM HEPES-NaOH, pH 7.5, 150 mM NaCl, 0.5 mM Spermidine (Sigma 85558-1G) supplemented with protease inhibitor cocktail (Sigma 4693132001) in 1.5ml tubes. Next, cells were resuspended in 1 ml of cold wash buffer, 20 µl of BioMagPlus Concanavalin A beads (Bangs laboratories BP531) were added and samples were mixed by rotation (4°C, 20 min). Samples were briefly spun at 100 g, placed on DynaMag™-2 Magnet (Thermo 12321D), and liquid was removed. Primary antibodies were diluted 1:100 in 250 µl of cold antibody buffer (20 mM HEPES-NaOH pH 7.5, 150 mM NaCl, 0.5 mM Spermidine, 2 mM EDTA, 0.1% digitonin (Millipore 300410-1GM) supplemented with protease inhibitor cocktails) and incubated with samples (4°C, overnight, with rotation). The following day, samples were washed once with 1 ml cold wash buffer. Protein A-MNase (pA-MN) was diluted 1:200 in 250 µl of cold digitonin buffer (20 mM HEPES-NaOH pH 7.5, 150 mM NaCl, 0.5 mM Spermidine, 0.1% digitonin supplemented with protease inhibitor cocktails) and added to samples (4°C, 1 h, with end-to-end rotation). Samples were washed twice with 1 ml of cold digitonin buffer, resuspended in 150 µl of cold digitonin buffer and placed on a pre-cooled metal block on ice for 5 min. pA-MN digestion was initiated by adding 3 µl of 0.1 M CaCl2 to samples, mixed by gently flicking tubes 20 times and samples placed back on metal block for 30 min. Digestion was stopped by adding 150 µl of 2 x stop buffer (340 mM NaCl, 20 mM EDTA, 4 mM EGTA, 0.02% Digitonin, 50 ug/ml RNase A (Thermo EN0531), 50 ug/ml Glycogen (Thermo R0561), and 4 pg/ml yeast heterologous spike-in DNA). Samples were incubated at 37°C for 10 min and then spun (16,000 g, 5 min, 4°C). Supernatant containing cleaved chromatin was transferred to a new tube, 3 µl of 10% SDS and 2.5 µl of 20 mg/ml proteinase K (Denville Scientific CB3210-5) were added and samples were incubated at 70°C for 10 min, followed by phenol:chloroform:isoamyl alcohol (Thermo 15593049) and chloroform (Sigma 288306) extraction. Supernatant containing DNA (∼300 ul) was transferred to new tubes pre-loaded with 20 ug of glycogen and then mixed with 750 µl of cold 100% ethanol for precipitation at -20°C overnight. Tubes were centrifuged at 20,000 g for 30 min at 4°C. DNA pellets were washed once with 1ml of cold 100% ethanol, air-dried, and stored at -20°C.

DNA libraries were prepared as previously described with slight modifications^94,116,117^. DNA pellets were dissolved in nuclease free H_2_O and library preparation performed using NEBNext Ultra II DNA Library Prep Kit (NEB E7645L). Adaptor was diluted to 1:25 for adaptor ligation. For samples labeled with CTCF, DNA was barcoded and amplified for 12 PCR cycles. For histone modification samples, adaptor-ligated DNA was first selected with 25 µl and second with 45 µl of AMPure XP beads, followed by PCR amplification (H3K4me3: 11 cycles, H3K9me3: 9 cycles, H3K27ac: 14 cycles, and H3K27me3: 10-12 cycles). All libraries were cleaned up using AMPure XP beads (Beckman Coulter A63881).

### CUT&RUN Antibodies

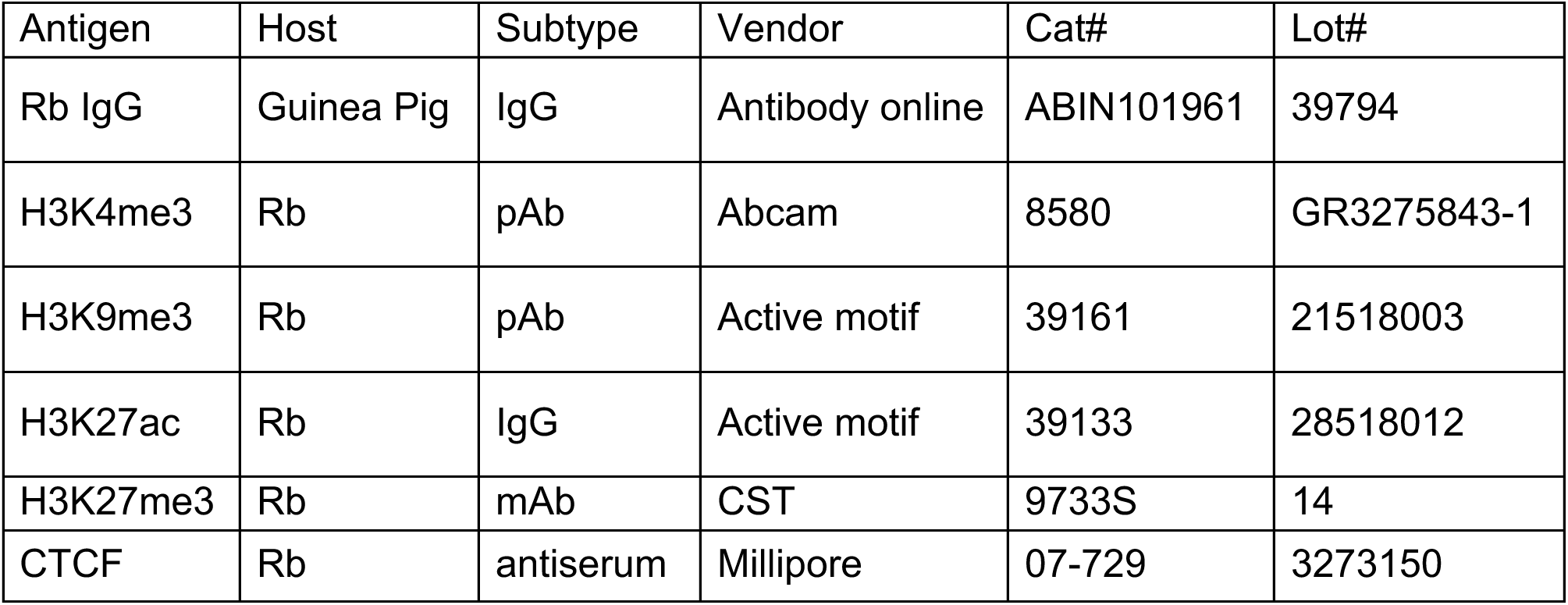

### RNA isolation

RNA-seq was performed as previously described with minor modifications^118^. 90,000 sorted cells were pelleted (600 g, 5 min, 4°C) and pellets washed once with cold PBS. Pellets were resuspended in 0.5 ml of TRIzol (Thermo 15596018) and stored at -80°C. Total RNA (∼30-120 ng) was extracted using RNA Clean & Concentrator-5 (ZYMO R1013) following manufacturer’s instruction and immediately followed by RNA library preparation. mRNA was isolated using NEBNext Poly(A) mRNA Magnetic Isolation Module (NEB E7490L). Libraries were prepared using NEBNext Ultra II Directional RNA Library Prep Kit (NEB E7760L) and following manufacturer’s instructions.

### Sequencing

Library quality was assessed using the Agilent 2100 Bioanalyzer (Agilent G2939BA) and libraries were quantified using a Qubit 2.0 fluorometer (Thermo Q32866) and by qPCR using NEBNext Library Quant Kit for Illumina (NEB E7630L) according to manufacturer’s instructions. Libraries were pooled at equal molarity and sequenced with NextSeq 500/550 High Output Kit (75 cycles) v2.5 kit (Illumina 20024906) on NextSeq 550 sequencing system (Illumina SY-415-1002). 20-30 million reads for each library were sequenced using paired-end sequencing (42:6:0:42).

### RNA-seq data processing and analysis

Paired-end reads were aligned and processed using STAR^119^ v2.7.1a with mm10 Gencode reference genome and default parameters. Paired-end read counts of genes were quantified by featureCounts using Gencode primary assembly annotation reference genome version vM24. Genes with raw reads greater than 5 were used for downstream analysis. Normalized read counts and differential analyses were generated using DESeq2^120^. Differentially expressed genes (DEGs) were identified with filters absolute fold change >1.5 and adjusted P-value <0.05. All pairwise DEGs across different cell types were combined and grouped into 7 clusters using the K-mean algorithm. Heatmap plots were generated using ComplexHeatmap^121,122^ packages. Other summary bar plots, violin plots, volcano plots were generated using R.

Pathway analysis of differentially expressed genes was performed using Metascape^123^. DESeq2^120^ normalized factors were used to normalize bam files. Normalized bigwig files were generated using bamCoverage^124^ with parameters -ignore chM -- minMappingQuality 5 -ignoreDuplicates -skipNAs and were visualized in UCSC genome browser and R package Gviz^125^.

### ATAC-seq data analysis and processing

ATAC-seq data were aligned and processed using Bowtie2^126^ v2.3.5 using mouse mm10 reference genome. Picard^127^ tools v1.96 was used to remove presumed PCR duplicates using the MarkDuplicates command. Bam files containing uniquely mapped reads were created using Samtools^128^ v1.1. Blacklist regions defined by ENCODE^129^, random chromosomes and mitochondria were removed, and filtered bam files were used for downstream analysis.

Union peaks were downloaded from GEO. Read per million (RPM/CPM) normalized bigwig files were created using deepTools bamCoverage^124,130^. Replicates were pooled together using wiggleTools and UCSC toolkit bedGraphToBigwig^131^, and tracks were imported and viewed using UCSC genome browser^132^.

### CUT&RUN data processing

FastQC^134^ v0.11.2 and MultiQC^135^ were used to check data quality. Reads were aligned to the mouse mm10 Gencode reference genome using Bowtie2^126^ v2.3.5, following parameters suggested by Skene et al.^115^ --local --very-sensitive-local --no-unal --no-mixed --no-discordant --phred33 -I 10 -X 700 -k1 -N1. Picard^127^ tools v1.96 was used to remove presumed PCR duplicates using MarkDuplicates command. Bam files containing uniquely mapped reads were created using Samtools^128,136^ v1.1. Fragments between 40-700 bp were kept. Blacklist regions defined by ENCODE, random chromosomes and mitochondria were removed, and filtered bam files were used for downstream analysis.

CUT&RUN signals were called using MACS^133^ v2.1 using the broadPeak setting with adjusted P-value cutoff 0.01. In broad histone modifications, customized parameters were adjusted for different hPTMs based on length of signals and sample background variation. Consensus peaks shown in at least two biological replicates were used, and merged conditional peaks with IgG peaks removal were finally used as a union peak list for downstream quantification. Venn diagrams were generated using the ChIPpeakAnno^137^ package findOverlapsOfPeaks() and makeVennDiagram() function.

Read counts were quantified across all samples based on union peak using featureCounts^138^, and validated using bedtools coverage. All sample read counts were normalized using DESeq2, and principal component analysis (PCA) plots of all replicates were generated using R function prcomp. Statistical significantly differential hPTMs analyses were performed using DESeq2. The histone modification regions with adjusted P-value < 0.05 and fold change > 1.5 were defined as significantly differential modification regions, and were quantified in bar plots using ggplots. The volcano plots were generated using R ggplot2. Approximate posterior estimation for GLM shrinkage method “apeglm”^139^ in DESeq2 was applied to H3K4me3 to alleviate its batch effects and final fold change calculation.

Binding motif enrichment of selected differential histone modification regions were identified using findMotifsGenome.pl from HOMER^140^ v4 using each corresponding union peaks as background and size as given with mask options.

For consistent visualization, DESeq2 normalization factors were used to adjust bam files to create normalized bigwig files using bamCoverage. Bigwig files of replicates were pooled together using WiggleTools^141^ mean setting. Tracks were loaded to UCSC genome browser and Gviz^125^ R package for visualization. Heatmaps and metaplots were generated using deepTools plotHeatmap^124,130^.

### CUT&RUN data analysis

#### Annotations -- Chromatin modification region annotation

Genes proximal to peaks (hPTMs) were annotated against mm10 genome using annotatePeaks.pl from HOMER^140^ v4, ChIPseeker^142^ with 10,000 base pairs (bp) flank regions, as well as GREAT^143^. Gene position information was extracted from the Gencode mm10 database, excluding pseudo genes or ambiguous undefined genes. Regions within 2,500bp of the TSS were defined as promoters. Annotation pie chart and bar chart of gene locations were generated based on filtered gene annotation.

For one-gene-one-peak mapping, the peak with maximum variations across different conditions representing the gene were selected. For one-gene-two-peak mapping, two peaks including the gene promoter and the non-promoter peak with maximum variation were used. For one-peak-multi-gene mapping, all genes annotated to target peaks were used. For hPTM patterns analysis and gene expression correlation, one-gene-one-peak mapping was used.

#### Taiji transcription factors analysis

Taiji^54^ analyses were performed using H3K27ac CUT&RUN bam data and RNA-seq read count data to predict key TFs. For comparison, the same analysis was performed using RNA-seq data and ATAC-seq from published papers^5,6^. The identified Taiji page rank scores of TFs were Z-score normalized across three conditions T_N_, T_MEM_ and T_EX_ to identify key TFs corresponding to each condition genomewide. Heatmaps were generated based on the page rank Z-scores, and plotted using ComplexHeatmap^121,122^.

Specific TF binding sites were identified using the MEME suite FIMO tool^144^. Venn diagrams of motif binding sites in H3K27ac and ATAC-seq were generated using ChIPpeakAnno^137^.

#### Super enhancer analysis

Promoter regions were defined as regions located within 2,500bp of TSS of each gene using the Gencode mm10 reference genome. Enhancers were defined as non-promoter regions with H3K27ac bound or ATAC-seq open accessibility. The enhancers between T_N_, T_MEM_ and T_EX_ were compared using ChIPpeakAnno^137^.

Super enhancers were identified using the stitching and rank ordering algorithm, ROSE^52,70^, using enhancers defined by H3K27ac and ATAC-seq, respectively. In brief, nearby enhancers were stitched together, ranked and plotted by signal enrichment levels. The enhancers with signals above tangent point (slope=1) were defined as super enhancers and the rest as typical enhancers. The SEs with only one peak stitched were further filtered out. SE were annotated to potential regulated genes using Homer^140^ and GREAT^143^, and further verified in the UCSC genome track. The SEs were ranked by signal intensity and plotted using R.

The comparison of SEs across T_N_, T_MEM_ and T_EX_ were performed as follows. First, an initial comparison and venn diagram were generated using ChIPpeakAnno^137^. Next, the SE signal intensities for overlapping and conditional-specific SEs were quantified for each cell type by summing the normalized reads of individual enhancers under the examined region. SEs were then filtered and refined: SEs with a fold change greater than 1.5 between any two conditions were classified as conditional-specific SEs, whereas other SEs were designated as shared respectively. An updated venn diagram was generated to reflect the refined SEs.

The enrichment signal intensity of SE and TE per condition was compared, and metaplots were generated using deepTools plotProfile^124,130^. Nearest genes were used to access gene expression differences between SE and TE, which were displayed in boxplot.

Comparison of SEs identified using H3K27ac and ATAC-Seq were generated using ChIPpeakAnno^137^, with hypergeometric P-values calculated for each pairwise comparison.

#### Bivalency analysis

Chromatin states were identified using chromHMM^75^ by separating promoters with non-promoters, respectively. Promoter regions were defined as regions located within 2,500bp of TSS of each gene and the rest of the regions were defined as non-promoter regions. ChromHMM was performed using consensus peaks of histone modifications H3K27ac, H3K27me3, H3K9me3 and ATAC with predefined 10 states, using concatenated mode with binarizing the peaks. The chromatin states coverage was quantified by base pairs. Initial states with non-low signals were further categorized into 4 major states, including I. active, II. poised, III. repressed and IV. repetitive states, based on the presence of histone modifications, and validated through heatmap plot of all marks in naive state.

Alluvial plots were generated to examine poised promoter dynamics from T_N_ to T_MEM_ and T_EX_ states. The significant changes were defined as histone changing with absolute fold change >1.5. The poised-to-activated promoters were defined as either gaining H3K4me3 or losing H3K27me3 or both, and poised-to-repressed promoters were defined as either losing H3K4me3 or gaining H3K27me3 or both. The identified dynamic regions were compared to significant DEGs.

The enriched motifs were identified using Homer^140^ v4 with parameters -size given and - mask, and all coding gene promoters were used as background for identifying all poised promoter motifs, while random genomes were used as background for identifying unique motifs enriched in the dynamic poised promoters. Pathway analyses were generated using Metascape on major chromatin states and dynamic promoters with consistent gene expression. Heatmaps were generated using R package ComplexHeatmap^122^. Examples of gene tracks were generated from UCSC genome tracks and Gviz^125^ R package. Summary bar plots and dot plots were generated using R ggplot2.

#### H3K9me3 analysis

Genome-wide H3K9me3 bound regions were split into broad and narrow peaks based on 15 kb cutoff. Peaks were annotated to nearest genes and genes within 250 kb.

Mouse (mm10) repeat database was downloaded from UCSC RepeatMasker tool^145,146^. Five runs of random background controls were generated using non-H3K9me3 bound genome regions with peak amount and width matching to T_EX_-enriched H3K9me3. The 5 random background regions were merged and used as one background control, average values were used for repeat coverage calculation.

Three levels of repeats, including repeat family, class and name, were used to calculate repeat coverage over peaks. Repeat covered regions were identified using bedtools^147^ intersect of query regions with the repeat database. Repeat coverage was calculated as base pairs covered by any type of repeats.

Comparisons of genome-wide H3K9me3 bound regions with CTCF binding sites and ATAC-seq open chromatin regions were performed using R package ChIPpeakAnno^137^ and tracks were generated using Gviz^125^.

## Acknowledgements

Protein A-MNase (batch 6) and yeast heterologous spike-in DNA were kindly provided by Dr. Steve Henikoff. The authors also thank Terri D. Bryson from Henikoff Laboratory for sharing the pA-MNase purification protocol. This work was supported by NIH grants AI155577, AI115712, AI117950, AI108545, AI082630, AI149680, HL145754 (to E.J.W.) and funding from Celgene (to S.L.B. and E.J.W.), Parker Institute for Cancer Immunotherapy (to E.J.W.), and The Mark Foundation (to E.J.W.). J.R.G. was supported by a Cancer Research Institute - Mark Foundation Fellowship. S.L.B. is supported by NIH grant CA078831. C.R.G is supported by NIH grant CA232466 and is a Rob Kugler American Cancer Society Postdoctoral fellow.

## Author contributions

E.J.W. and S.L.B. conceived the project. A.E.B., Z.C., Z.Z., P.A.AG., E.J.W. and S.L.B. designed the experiments. A.E.B., Z.C., Z.Z., P.A.AG., P.S. and S.C. performed experiments. H.H. analyzed data and prepared figures with help from A.E.B., Z.Z., C.R.G and K.A.A., K.M.G., L.W., G.D., S.M., J.R.G. and J.S. consulted on data analysis. H.H., A.E.B., Z.Z., C.R.G, K.A.A., S.L.B. and E.J.W. wrote the manuscript. All authors reviewed the manuscript.

## Ethical approval

This study is reported in accordance with ARRIVE guidelines.

## Data availability

RNA-seq and CUT&RUN data generated in this study will be deposited in the National Center for Biotechnology Information Gene Expression Omnibus and made available after publication. ATAC-seq data used in this study is from GSE86797.

