## Supplemental Figures for "Deciphering the role of histone modifications in memory and exhausted CD8 T cells"

#### SUPPLEMENTARY FIGURE LEGENDS

**Figure S1. Major changes in hPTM occur as $T_N$ differentiate into $T_{MEM}$ or $T_{EX}$**

(**a**) Gating strategy for P14 cell isolation from acute LCMV Armstrong infection ($T_{MEM}$).

(**b**) Post-isolation purity for $T_{MEM}$.

(**c**) Flow cytometry plot showing KLRG1 versus CD127 expression on isolated $T_{MEM}$.

(**d**) Gating strategy for P14 cell isolation from chronic LCMV clone13 infection ($T_{EX}$).

(**e**) Post-isolation purity for $T_{EX}$.

(**f**) Flow cytometry plot showing PD-1 expression for isolated $T_{EX}$.

(**g**) Pairwise comparisons between CD8 T cell subsets showing percent and number of differentially hPTMs for each histone modification, with adjusted P-value < 0.05 and absolute fold change > 1.5. Total number of union peaks (N) indicated.

(**h**) Correlation of change in RNA expression between $T_{MEM}$ and $T_N$ and the change in H3K27ac (left), and H3K27me3 (right). R and associated P-value represent Pearson correlation.

(**i**) Correlation of change in RNA expression between $T_{EX}$ and $T_N$ and the change in H3K27ac (left), and H3K27me3 (right). R and associated P-value represent Pearson correlation.

(**j**) Genome track showing RNA-seq, ATAC-seq and hPTM data. Differentially modified regions between $T_{MEM}$ and $T_{EX}$ for each hPTM are highlighted in black boxes.

(**k**) Volcano plots comparing hPTMs between $T_{MEM}$ and $T_{EX}$. Genes with increased levels of indicated hPTM in $T_{EX}$ are right (red). Genes with increased levels of indicated hPTM in $T_{MEM}$ are left (blue). Select genes are labeled.

**Figure S2. Predicted NUR77 binding motifs increase in chromatin accessibility in $T_{EX}$ without concurrent H3K27ac changes**

(**a**) Heatmap of normalized Taiji PageRank scores calculated using RNA-seq and ATAC-seq data.

(**b**) Venn diagram comparing number of NUR77 motifs in regions with increased H3K27ac in $T_{EX}$ to regions with increased chromatin accessibility (ATAC) in $T_{EX}$.

(**c**) Heatmap showing DEGs between $T_{MEM}$ and $T_{EX}$ associated with regions with increased chromatin accessibility in $T_{EX}$ that contain NUR77 motifs.

(**d**) Bar graph showing cell type expression of DEGs associated with NUR77 motifs in regions with increased chromatin accessibility without H3K27ac changes in $T_{EX}$.

(**e**) Genome tracks highlighting NUR77 motifs in regions with increased chromatin accessibility in $T_{EX}$ without changing H3K27ac. Differentially modified regions for H3K27ac and open chromatin are highlighted in boxes under tracks. Predicted NUR77 binding sites are shown in red.

(**f**) Comparison of hPTMs between $T_{EX}$ and $T_{MEM}$ for TFs with increased expression in $T_{MEM}$. Top six most frequent groups plotted.

**Figure S3. H3K27ac and chromatin accessibility identify shared and distinct SEs**

(**a**) Venn diagram showing cell-type specificity of total enhancers.

(**b**) Metaplot of enrichment signals for SEs compared to TEs in $T_N$, $T_{MEM}$ and $T_{EX}$.

(**c**) Gene expression associated with SEs compared to TEs in $T_N$, $T_{MEM}$ and $T_{EX}$.

(**d**) Distribution of ATAC-seq signal across stitched enhancer regions in $T_N$, $T_{MEM}$ and $T_{EX}$. Top 10 putative SEs labeled. N represents the number of putative SEs identified from each cell type. Stitched enhancers above horizontal dashed line are associated with putative SEs.

(**e**) Venn diagram comparing SEs identified by ATAC-seq and H3K27ac for $T_N$, $T_{MEM}$ and $T_{EX}$. P value represents a hypergeometric test between each comparison.

(**f**) Genome tracks showing RNA-seq, ATAC-seq and H3K27ac data. Putative SEs identified in ATAC-seq but not H3K27ac data are highlighted in black boxes below tracks. Individual enhancers within SE regions are highlighted in grey. Promoter regions (±2,500bp of TSS) are indicated in red.

(**g**) Pathway analysis comparing DEGs (from Fig. 3e) potentially regulated by SEs identified by either H3K27ac or ATAC.

**Figure S4. Genes poised in $T_N$ are actively repressed in $T_{MEM}$ and $T_{EX}$**

(**a**) Heatmap showing characterization of chromatin states at promoters.

(**b**) Signal intensity heatmap showing hPTM density at promoters associated with each chromatin state.

(**c**) Bar plot showing frequencies of four promoter chromatin states in $T_N$ and associated gene expression. Genes in the lowest 25 percentile of gene expression were defined as low expression genes.

(**d**) Top10 GO pathway terms for four major promoter chromatin states in $T_N$.

(**e**) GO analysis comparing poised-to-activated promoters in $T_{EX}$ versus $T_{MEM}$.

(**f**) Genome tracks of RNA-seq, H3K4me3 and H3K27me3 data showing chromatin changes at the *Id2* promoter.

(**g**) Boxplot quantifying log2-scaled normalized read count (RC) of H3K4me3 and H3K27me3 for promoters that switched from poised-to-repressed in $T_{MEM}$.

(**h**) Heatmap of normalized RC of H3K4me3 and H3K27me3 for promoters that switched from poised-to-repressed in $T_{MEM}$ (top) and $T_{EX}$ (bottom), with selected genes labeled.

(**i**) Genome tracks of RNA-seq, H3K4me3 and H3K27me3 data showing example promoters that switched from poised-to-repressed in $T_{MEM}$.

(**j**) Boxplot quantifying log2-scaled normalized RC of H3K4me3 and H3K27me3 for promoters that switched from poised-to-repressed in $T_{EX}$.

(**k**) Genome tracks of RNA-seq, H3K4me3 and H3K27me3 data showing example promoters that switched from poised-to-repressed in $T_{EX}$.

**Figure S5. $T_{EX}$-enriched H3K9me3 peaks associated with increased expression of genes involved in diverse cellular processes**

(**a**) Signal intensity heatmap of union H3K9me3 regions between all three CD8 T cell populations.

(**b**) Percentage of genome base pair coverage for broad (>=15 kb) and narrow (<15 kb) H3K9me3 peaks. N.s. = non-significantly different between $T_{EX}$ and $T_{MEM}$.

**(c)** Genomic locations of $T_{EX}$-enriched H3K9me3 narrow peaks.

**(d)** Bar chart showing percentage of repeat and non-repeat coverage for $T_{MEM}$-enriched, n.s. and $T_{EX}$-enriched H3K9me3 narrow and broad peaks.

**(e)** Pie chart showing repeat element family coverage for $T_{EM}$-enriched, n.s. and $T_{EX}$-enriched H3K9me3 narrow and broad peaks.

**(f)** Genome track showing example genes associated with $T_{EX}$-enriched narrow H3K9me3 peaks. The $T_{EX}$-enriched narrow H3K9me3 peaks with predicted CTCF binding sites highlighted in grey.

**(g)** Venn diagram showing the overlaps between ATAC-seq open chromatin regions and $T_{EX}$-enriched H3K9me3 narrow peaks.

**(h)** Gene ontology pathways for genes with $T_{EX}$-enriched narrow H3K9me3, with and without ATAC-seq open chromatin in $T_{EX}$.

**(i)** Example gene track displaying loci with $T_{EX}$-enriched H3K9me3 narrow peaks and ATAC-seq open chromatin in $T_{EX}$.

**(j)** Violin plot of log2 fold change in RNA expression between $T_{MEM}$ and $T_{EX}$ of genes near differentially modified broad and narrow H3K9me3 peaks.

**(k)** Pie chart showing changes in RNA expression between $T_{MEM}$ and $T_{EX}$ of genes near $T_{MEM}$-enriched narrow H3K9me3 peaks.

**(l)** Bar chart showing percentage of genes upregulated in $T_{EX}$ versus $T_{MEM}$ located close to $T_{EX}$-enriched narrow peaks. Three different base pair window sizes around DEGs were used to identify nearby H3K9me3 peaks.

**(m)** Gene ontology pathways for DEGs with increased or decreased expression in $T_{EX}$ compared to $T_{MEM}$ associated with nearby $T_{EX}$-enriched narrow H3K9me3 peaks.

### Figure S1

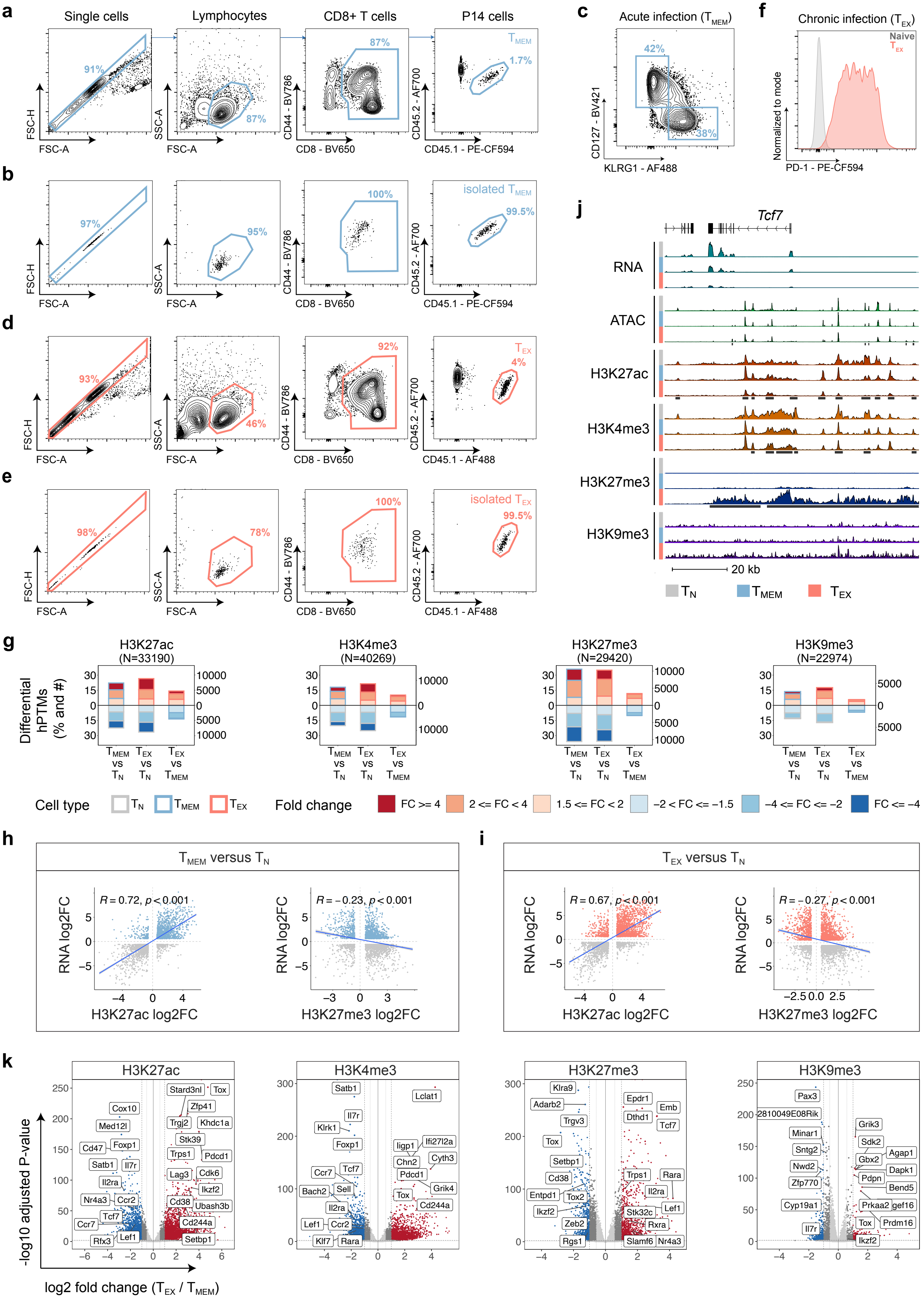

a
Single cells
Lymphocytes
CD8+ T cells
P14 cells
91%
87%
87%
1.7%
TMEM
FSC-H
FSC-A
SSC-A
CD44 - BV786
CD8 - BV650
CD45.2 - AF700
CD45.1 - PE-CF594
b
97%
95%
100%
99.5%
isolated TMEM
c
Acute infection (TMEM)
42%
38%
CD127 - BV421
KLRG1 - AF488
f
Chronic infection (TEX)
Naive
TEX
Normalized to mode
PD-1 - PE-CF594
d
93%
46%
92%
4%
TEX
CD45.1 - AF488
e
98%
78%
100%
99.5%
isolated TEX
j
Tcf7
RNA
ATAC
H3K27ac
H3K4me3
H3K27me3
H3K9me3
20 kb
TN
TMEM
TEX
g
H3K27ac (N=33190)
H3K4me3 (N=40269)
H3K27me3 (N=29420)
H3K9me3 (N=22974)
Differential hPTMs (% and #)
TMEM vs TN
TEX vs TN
TEX vs TMEM
Cell type
Fold change
FC >= 4
2 <= FC < 4
1.5 <= FC < 2
−2 < FC <= −1.5
−4 <= FC <= −2
FC <= −4
h
TMEM versus TN
R = 0.72, p < 0.001
R = −0.23, p < 0.001
RNA log2FC
H3K27ac log2FC
H3K27me3 log2FC
i
TEX versus TN
R = 0.67, p < 0.001
R = −0.27, p < 0.001
k
H3K27ac
H3K4me3
H3K27me3
H3K9me3
-log10 adjusted P-value
log2 fold change (TEX / TMEM)
Stard3nl
Tox
Cox10
Zfp41
Med12l
Trgj2
Khdc1a
Cd47
Foxp1
Stk39
Trps1
Pdcd1
Satb1
Il7r
Il2ra
Lag3
Cdk6
Ikzf2
Nr4a3
Ccr2
Cd38
Ubash3b
Tcf7
Cd244a
Ccr7
Setbp1
Rfx3
Lef1
Lclat1
Klrk1
Iigp1
Ifi27l2a
Chn2
Cyth3
Pdcd1
Grik4
Bach2
Sell
Klf7
Rara
Klra9
Epdr1
Adarb2
Emb
Dthd1
Trgv3
Entpd1
Tox2
Stk32c
Zeb2
Rxra
Rgs1
Slamf6
Nr4a3
Pax3
2810049E08Rik
Grik3
Minar1
Sdk2
Sntg2
Agap1
Gbx2
Dapk1
Nwd2
Pdpn
Zfp770
Bend5
Cyp19a1
Prkaa2
gef16
Prdm16
Ikzf2

#### Figure S2

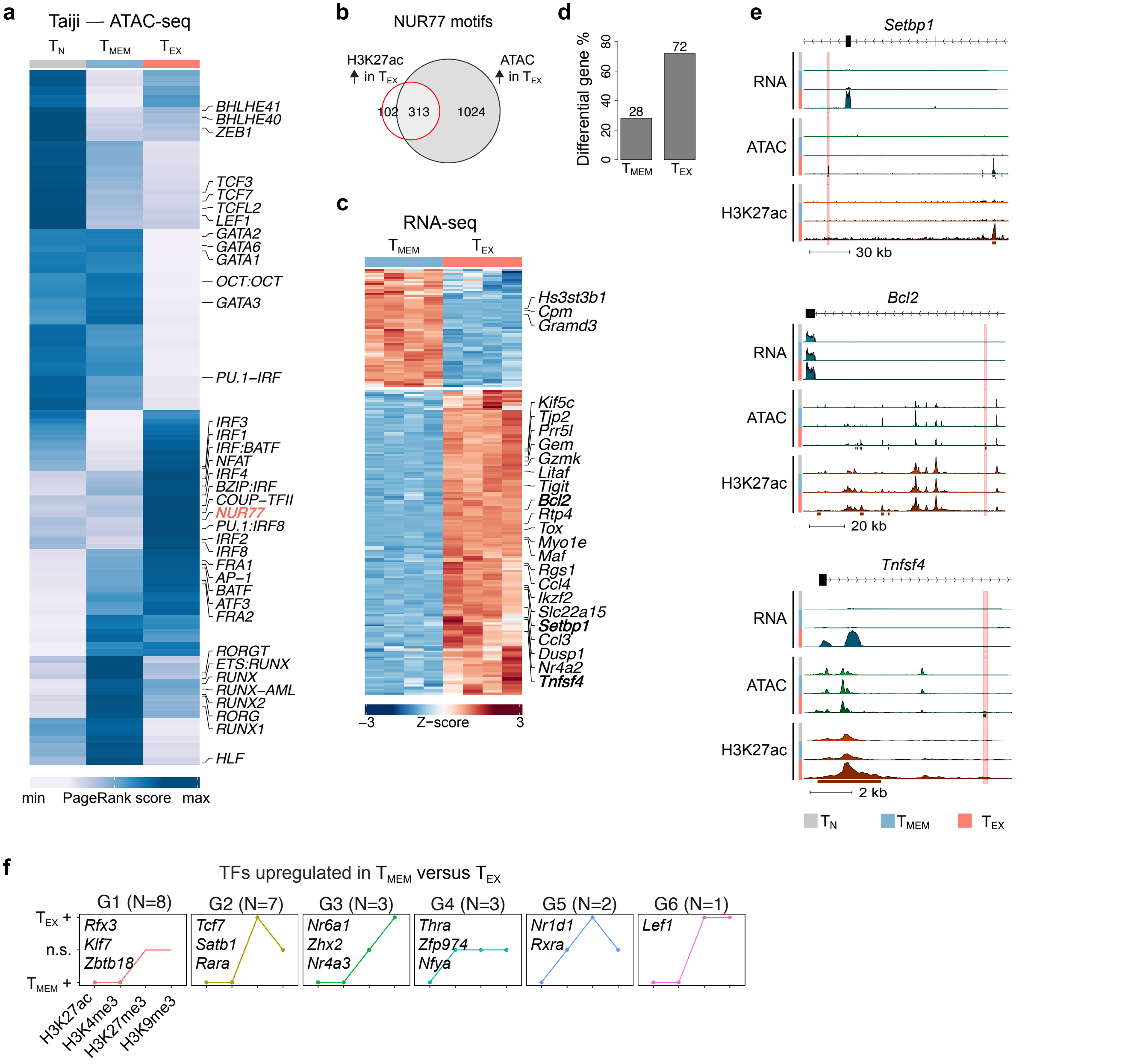

a
Taiji — ATAC-seq
$T_N$
$T_{MEM}$
$T_{EX}$
BHLHE41
BHLHE40
ZEB1
TCF3
TCF7
TCFL2
LEF1
GATA2
GATA6
GATA1
OCT:OCT
GATA3
PU.1–IRF
IRF3
IRF1
IRF:BATF
NFAT
IRF4
BZIP:IRF
COUP–TFII
NUR77
PU.1:IRF8
IRF2
IRF8
FRA1
AP–1
BATF
ATF3
FRA2
RORGT
ETS:RUNX
RUNX
RUNX–AML
RUNX2
RORG
RUNX1
HLF
min
PageRank score
max
b
NUR77 motifs
H3K27ac in $T_{EX}$
ATAC in $T_{EX}$
102
313
1024
c
RNA-seq
$T_{MEM}$
$T_{EX}$
Hs3st3b1
Cpm
Gramd3
Kif5c
Tjp2
Prr5l
Gem
Gzmk
Litaf
Tigit
Bcl2
Rtp4
Tox
Myo1e
Maf
Rgs1
Ccl4
Ikzf2
Slc22a15
Setbp1
Ccl3
Dusp1
Nr4a2
Tnfsf4
–3
Z–score
3
d
Differential gene %
0
20
40
60
80
28
72
$T_{MEM}$
$T_{EX}$
e
Setbp1
RNA
ATAC
H3K27ac
30 kb
Bcl2
RNA
ATAC
H3K27ac
20 kb
Tnfsf4
RNA
ATAC
H3K27ac
2 kb
$T_N$
$T_{MEM}$
$T_{EX}$
f
TFs upregulated in $T_{MEM}$ versus $T_{EX}$
G1 (N=8)
G2 (N=7)
G3 (N=3)
G4 (N=3)
G5 (N=2)
G6 (N=1)
$T_{EX}$ +
n.s.
$T_{MEM}$ +
Rfx3
Klf7
Zbtb18
Tcf7
Satb1
Rara
Nr6a1
Zhx2
Nr4a3
Thra
Zfp974
Nfya
Nr1d1
Rxra
Lef1
H3K27ac
H3K4me3
H3K27me3
H3K9me3

Figure S3

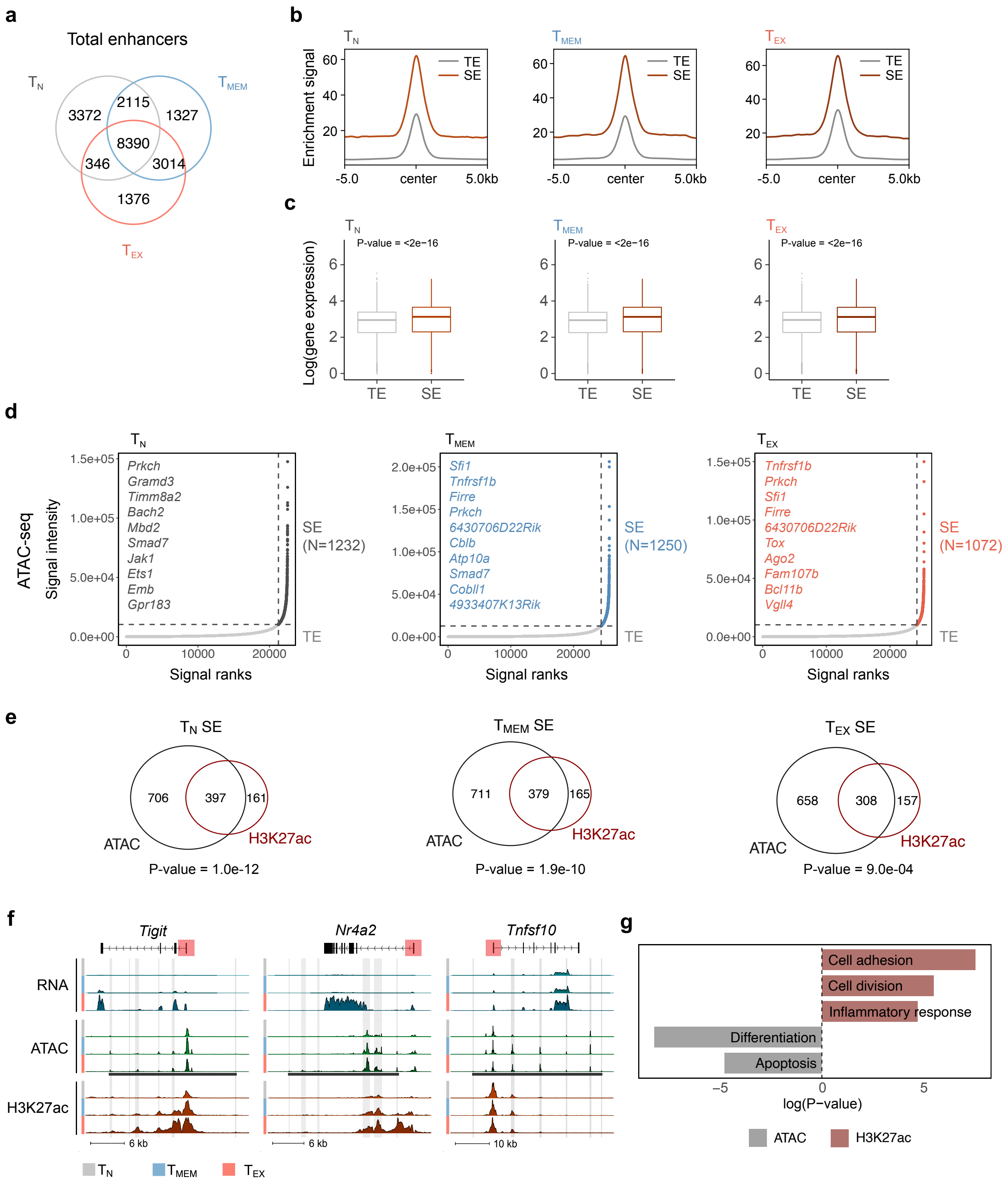
a
Total enhancers
T N
T MEM
T EX
3372
2115
1327
8390
346
3014
1376
b
Enrichment signal
TE
SE
-5.0
center
5.0kb
c
Log(gene expression)
P-value = <2e−16
TE
SE
d
ATAC-seq
Signal intensity
Signal ranks
Prkch
Gramd3
Timm8a2
Bach2
Mbd2
Smad7
Jak1
Ets1
Emb
Gpr183
SE (N=1232)
Sfi1
Tnfrsf1b
Firre
Prkch
6430706D22Rik
Cblb
Atp10a
Smad7
Cobll1
4933407K13Rik
SE (N=1250)
Tnfrsf1b
Prkch
Sfi1
Firre
6430706D22Rik
Tox
Ago2
Fam107b
Bcl11b
Vgll4
SE (N=1072)
e
T N SE
T MEM SE
T EX SE
706
397
161
711
379
165
658
308
157
ATAC
H3K27ac
P-value = 1.0e-12
P-value = 1.9e-10
P-value = 9.0e-04
f
Tigit
Nr4a2
Tnfsf10
RNA
ATAC
H3K27ac
6 kb
10 kb
g
Cell adhesion
Cell division
Inflammatory response
Differentiation
Apoptosis
log(P-value)
ATAC
H3K27ac

#### Figure S4

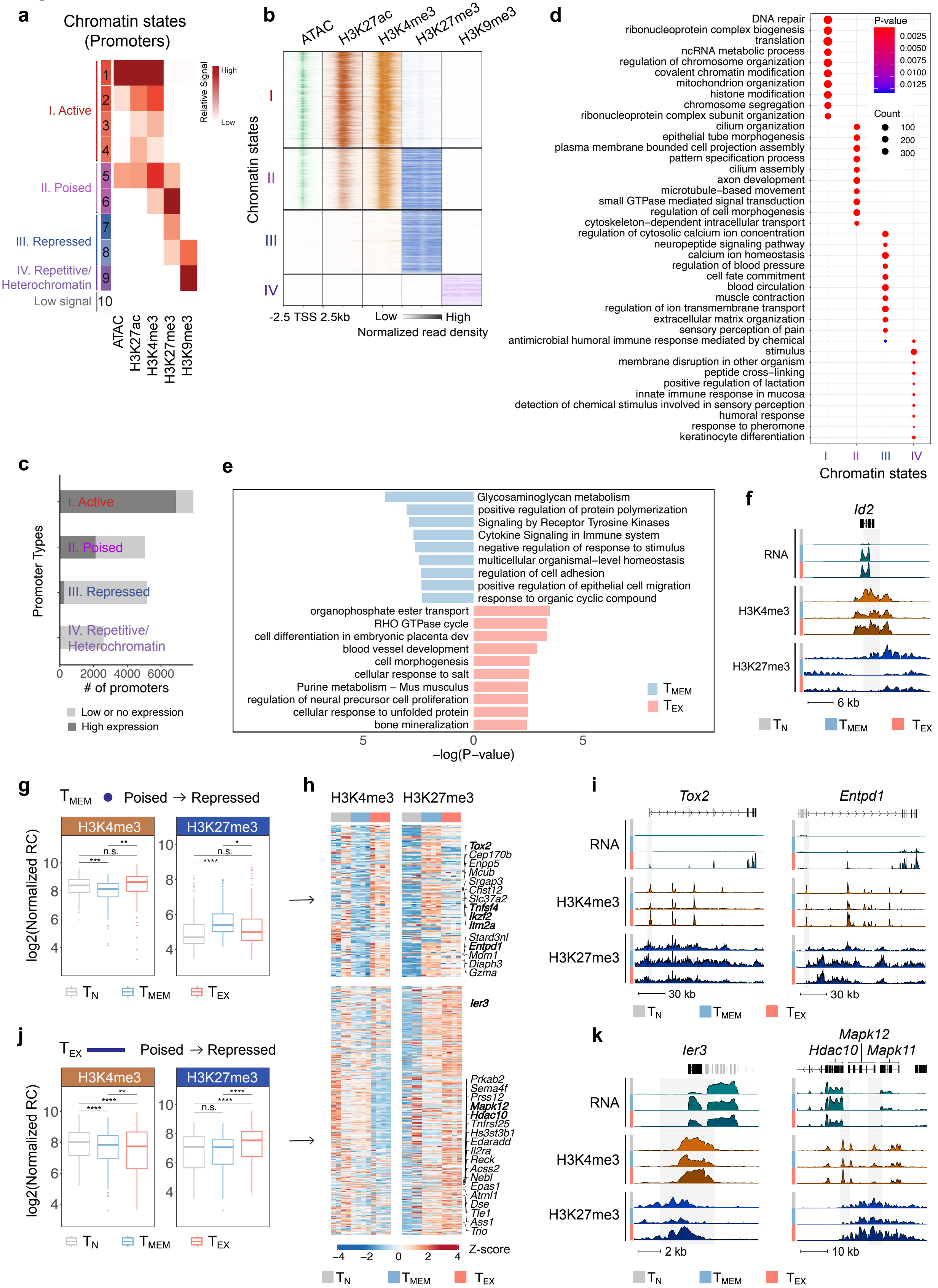

a
Chromatin states (Promoters)
I. Active
II. Poised
III. Repressed
IV. Repetitive/ Heterochromatin
Low signal
Relative Signal
High
Low
ATAC
H3K27ac
H3K4me3
H3K27me3
H3K9me3
b
Chromatin states
-2.5 TSS 2.5kb
Low High
Normalized read density
c
Promoter Types
### of promoters
Low or no expression
High expression
d
DNA repair
ribonucleoprotein complex biogenesis
translation
ncRNA metabolic process
regulation of chromosome organization
covalent chromatin modification
mitochondrion organization
histone modification
chromosome segregation
ribonucleoprotein complex subunit organization
cilium organization
epithelial tube morphogenesis
plasma membrane bounded cell projection assembly
pattern specification process
cilium assembly
axon development
microtubule−based movement
small GTPase mediated signal transduction
regulation of cell morphogenesis
cytoskeleton−dependent intracellular transport
regulation of cytosolic calcium ion concentration
neuropeptide signaling pathway
calcium ion homeostasis
regulation of blood pressure
cell fate commitment
blood circulation
muscle contraction
regulation of ion transmembrane transport
extracellular matrix organization
sensory perception of pain
antimicrobial humoral immune response mediated by chemical stimulus
membrane disruption in other organism
peptide cross−linking
positive regulation of lactation
innate immune response in mucosa
detection of chemical stimulus involved in sensory perception
humoral response
response to pheromone
keratinocyte differentiation
P-value
Count
Chromatin states
e
Glycosaminoglycan metabolism
positive regulation of protein polymerization
Signaling by Receptor Tyrosine Kinases
Cytokine Signaling in Immune system
negative regulation of response to stimulus
multicellular organismal−level homeostasis
regulation of cell adhesion
positive regulation of epithelial cell migration
response to organic cyclic compound
organophosphate ester transport
RHO GTPase cycle
cell differentiation in embryonic placenta dev
blood vessel development
cell morphogenesis
cellular response to salt
Purine metabolism – Mus musculus
regulation of neural precursor cell proliferation
cellular response to unfolded protein
bone mineralization
−log(P−value)
f
Id2
RNA
H3K4me3
H3K27me3
6 kb
g
$T_{MEM}$ Poised → Repressed
log2(Normalized RC)
h
Z-score
i
Tox2
Entpd1
30 kb
j
$T_{EX}$ Poised → Repressed
k
Ier3
Mapk12
Hdac10
Mapk11
2 kb
10 kb
$T_N$ $T_{MEM}$ $T_{EX}$

#### Figure S5

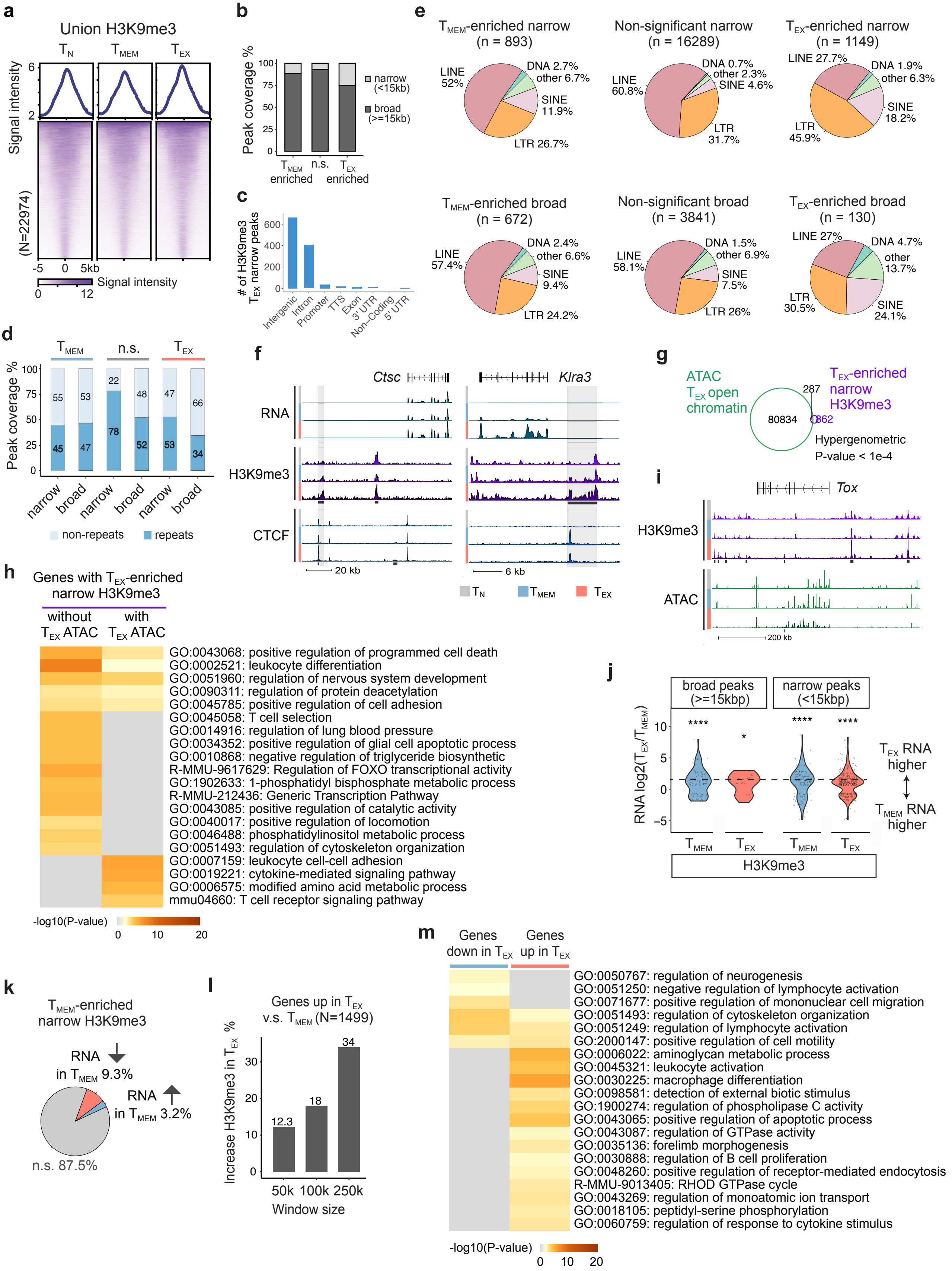

a
Union H3K9me3
T_N T_MEM T_EX
Signal intensity
(N=22974)
-5 0 5kb
Signal intensity
0 12
b
Peak coverage %
narrow (<15kb)
broad (>=15kb)
T_MEM enriched
n.s.
T_EX enriched
c
### of H3K9me3 T_EX narrow peaks
Intergenic
Intron
Promoter
TTS
Exon
3' UTR
Non-Coding
5' UTR
d
T_MEM n.s. T_EX
Peak coverage %
55 53 22 48 47 66
45 47 78 52 53 34
narrow broad narrow broad narrow broad
non-repeats
repeats
e
T_MEM-enriched narrow (n = 893)
LINE 52%
DNA 2.7%
other 6.7%
SINE 11.9%
LTR 26.7%
Non-significant narrow (n = 16289)
LINE 60.8%
DNA 0.7%
other 2.3%
SINE 4.6%
LTR 31.7%
T_EX-enriched narrow (n = 1149)
LINE 27.7%
DNA 1.9%
other 6.3%
SINE 18.2%
LTR 45.9%
T_MEM-enriched broad (n = 672)
LINE 57.4%
DNA 2.4%
other 6.6%
SINE 9.4%
LTR 24.2%
Non-significant broad (n = 3841)
LINE 58.1%
DNA 1.5%
other 6.9%
SINE 7.5%
LTR 26%
T_EX-enriched broad (n = 130)
LINE 27%
DNA 4.7%
other 13.7%
SINE 24.1%
LTR 30.5%
f
Ctsc
Klra3
RNA
H3K9me3
CTCF
20 kb
6 kb
T_N T_MEM T_EX
g
ATAC T_EX open chromatin
T_EX-enriched narrow H3K9me3
80834
287
862
Hypergenometric P-value < 1e-4
h
Genes with T_EX-enriched narrow H3K9me3
without T_EX ATAC
with T_EX ATAC
GO:0043068: positive regulation of programmed cell death
GO:0002521: leukocyte differentiation
GO:0051960: regulation of nervous system development
GO:0090311: regulation of protein deacetylation
GO:0045785: positive regulation of cell adhesion
GO:0045058: T cell selection
GO:0014916: regulation of lung blood pressure
GO:0034352: positive regulation of glial cell apoptotic process
GO:0010868: negative regulation of triglyceride biosynthetic
R-MMU-9617629: Regulation of FOXO transcriptional activity
GO:1902633: 1-phosphatidyl bisphosphate metabolic process
R-MMU-212436: Generic Transcription Pathway
GO:0043085: positive regulation of catalytic activity
GO:0040017: positive regulation of locomotion
GO:0046488: phosphatidylinositol metabolic process
GO:0051493: regulation of cytoskeleton organization
GO:0007159: leukocyte cell-cell adhesion
GO:0019221: cytokine-mediated signaling pathway
GO:0006575: modified amino acid metabolic process
mmu04660: T cell receptor signaling pathway
-log10(P-value)
0 10 20
i
Tox
H3K9me3
ATAC
200 kb
j
broad peaks (>=15kbp)
narrow peaks (<15kbp)
RNA log2(T_EX/T_MEM)
**** * **** ****
T_EX RNA higher
T_MEM RNA higher
T_MEM T_EX T_MEM T_EX
H3K9me3
k
T_MEM-enriched narrow H3K9me3
RNA in T_MEM 9.3%
RNA in T_MEM 3.2%
n.s. 87.5%
l
Genes up in T_EX v.s. T_MEM (N=1499)
Increase H3K9me3 in T_EX %
12.3 18 34
50k 100k 250k
Window size
m
Genes down in T_EX
Genes up in T_EX
GO:0050767: regulation of neurogenesis
GO:0051250: negative regulation of lymphocyte activation
GO:0071677: positive regulation of mononuclear cell migration
GO:0051493: regulation of cytoskeleton organization
GO:0051249: regulation of lymphocyte activation
GO:2000147: positive regulation of cell motility
GO:0006022: aminoglycan metabolic process
GO:0045321: leukocyte activation
GO:0030225: macrophage differentiation
GO:0098581: detection of external biotic stimulus
GO:1900274: regulation of phospholipase C activity
GO:0043065: positive regulation of apoptotic process
GO:0043087: regulation of GTPase activity
GO:0035136: forelimb morphogenesis
GO:0030888: regulation of B cell proliferation
GO:0048260: positive regulation of receptor-mediated endocytosis
R-MMU-9013405: RHOD GTPase cycle
GO:0043269: regulation of monoatomic ion transport
GO:0018105: peptidyl-serine phosphorylation
GO:0060759: regulation of response to cytokine stimulus
-log10(P-value)
0 10 20
